# Microbial strategies for survival in the glass sponge *Vazella pourtalesii*

**DOI:** 10.1101/2020.05.28.122663

**Authors:** Kristina Bayer, Kathrin Busch, Ellen Kenchington, Lindsay Beazley, Sören Franzenburg, Jan Michels, Ute Hentschel, Beate M. Slaby

## Abstract

Few studies have thus far explored the microbiomes of glass sponges (Hexactinellida). The present study seeks to elucidate the composition of the microbiota associated with the glass sponge *Vazella pourtalesii* and the functional strategies of the main symbionts. We combined microscopic approaches with metagenome-guided microbial genome reconstruction and amplicon community profiling towards this goal. Microscopic imaging revealed that the host and microbial cells appeared within dense biomass patches that are presumably syncytial tissue aggregates. Based on abundances in amplicon libraries and metagenomic data, SAR324 bacteria, Crenarchaeota, Patescibacteria and Nanoarchaeota were identified as abundant members of the *V. pourtalesii* microbiome and their genomic potentials were thus analyzed in detail. A general pattern emerged in that the *V. pourtalesii* symbionts had very small genome sizes in the range of 0.5-2.2 Mb and low GC contents, even below those of seawater relatives. Based on functional analyses of metagenome-assembled genomes (MAGs), we propose two major microbial strategies: the “givers”, namely Crenarchaeota and SAR324, heterotrophs and facultative anaerobes, produce and partly secrete all required amino acids and vitamins. The “takers”, Nanoarchaeota and Patescibacteria, are anaerobes with reduced genomes that tap into the microbial community for resources, e.g., lipids and DNA, likely using pili-like structures. We posit that the existence of microbial cells in sponge syncytia together with the low-oxygen conditions in the seawater environment are factors that shape the unique compositional and functional properties of the microbial community associated with *V. pourtalesii*.

**Importance:** We investigated the microbial community of *V. pourtalesii* that forms globally unique, monospecific sponge grounds under low-oxygen conditions on the Scotian Shelf, where it plays a key role for its vulnerable ecosystem. The microbial community was found to be concentrated within biomass patches and is dominated by small cells (<1 μm). MAG analyses showed consistently small genome sizes and low GC contents, which is unusual in comparison to known sponge symbionts. These properties as well as the (facultatively) anaerobic metabolism and a high degree of interdependence between the dominant symbionts regarding amino acid and vitamin synthesis are likely adaptations to the unique conditions within the syncytial tissue of their hexactinellid host and the low-oxygen environment.

## Introduction

The fossil record shows that sponges (Porifera) have been essential members of reef communities in various phases of Earth’s history, and even built biohermal reefs in the mid-Jurassic to early-Cretaceous (1). Today, extensive sponge aggregations, also known as ‘sponge grounds’, are found throughout the World’s oceans from temperate to arctic regions along shelves, on ridges, and seamounts (2). They can be mono-to multispecific with a single or various sponge species dominating the benthic community, respectively. In sponge ground ecosystems, these basal animals play a crucial role in the provision of habitat, adding structural complexity to the environment and thereby attracting other organisms, ultimately causing an enhancement of local biodiversity (3–5).

Studies on demosponges have shown that they harbor distinct and diverse microbial communities that interact with each other, their host and the environment in various ways (6, 7). The microbial consortia of sponges are represented by diverse bacterial and archaeal communities with ≥63 prokaryotic phyla having been found in sponges so far (6, 8). These sponge microbiomes display host species-specific patterns that are distinctly different from those of seawater in terms of richness, diversity and community composition. Microbial symbionts contribute to holobiont metabolism (e.g., via nitrogen cycling and vitamin production) and defense (e.g., via secondary metabolite production), (reviewed in (7)). Sponges and their associated microbial communities (hereafter termed “holobionts”) further contribute to fundamental biogeochemical cycles like nitrogen, phosphorous, and dissolved organic matter in the ecosystem, but the relative contribution of microbial symbionts remains mostly unresolved (1, 7, 9, 10).

Sponges of the class Hexactinellida (glass sponges) are largely present and abundant in the mesopelagic realm below 400 feet. They can form extensive reefs of biohermal character and can dominate the sponge ground ecosystems (1, 11). Glass sponges are characterized by a skeleton of siliceous spicules that is six-rayed symmetrical with square axial proteinaceous filament (12). Much of the body is composed of syncytial tissue which represents extensive and continous regions of multinucleated cytoplasm (12, 13). Also nutrients are transported via the cytoplasmic streams of these trabecular syncytia (12). Some discrete cell types exist, including choanocytes and the pluripotent archaeocytes that are likely non-motile and thus not involved in nutrient transport in Hexactinellida (12). While the microbial symbiont diversity and functions are well studied in Demospongiae, much less is known about the presence and function of microbes in glass sponges. In fact, microorganisms have rarely been seen in glass sponges (12). A recent study of South Pacific sponge microbial communities has however shown that general patterns seen previously in shallow-water sponge microbiomes, such as host-specificity and LMA-HMA dichotomy, are generally applicable for these deep-sea sponge microbiomes as well, including those of glass sponges (14). Another study underlined the importance of ammonia-oxidizing archaea (family Nitrosopumilaceae, phylum Thaumarchaeota) in the deep-sea hexactinellid *Lophophysema eversa* using metagenomic data (15).

Here we investigate the microbial community of *Vazella pourtalesii* (16), a glass sponge (class Hexactinellida) that forms globally unique, monospecific sponge grounds on the Scotian Shelf off Nova Scotia, Canada. This ecosystem is characterized by relatively warm and nutrient rich water with low oxygen concentrations (1, 17, 18). While there have been a number of studies on the distribution, biomarkers, and possible functional roles of *V. pourtalesii* in the ecosystem (1, 17, 19), little has been published to date on its associated microbiota (20). According to phylogenetic and fossil studies, sponges (including glass sponges) originate from Neoproterozoic times when oxygen was limited (21). Moreover, laboratory experiments have shown that sponges can cope with low oxygen levels for extended periods of time (22–24). Due to the observed low-oxygen conditions at the sampling location, we explored whether the *V. pourtalesii* microbiome contains compositional as well as functional adaptations to the low oxygen environment. Microscopy, metagenome-guided microbial genome reconstruction and amplicon community profiling were employed towards this goal.

## Results

### Site description

On the Scotian Shelf off Nova Scotia, eastern Canada, highest densities of *V. pourtalesii* were observed and/or predicted in the Emerald Basin and the Sambro Bank areas (Figure 1A). The water column of this region has a characteristic vertical structure with water masses of different temperatures and salinities gradually mixing and creating a distinct temperature-salinity profile (Figure 1B). Main water masses influencing the sampling sites are from surface to deep sea: Cabot Strait Subsurface Water (CBS), Inshore Labrador Current (InLC), Cold Intermediate Layer of Cabot Strait Subsurface Water (CBS-CIL), Labrador Slope Water (LSW), Warm Slope Water (WSW). All *V. pourtalesii* samples of this study originate from a relatively warm (>10 °C) and nutrient rich water mass called Warm Slope Water (WSW) which originates from the Gulf Stream (18). Relatively low oxygen concentrations (<4 mL/L) were measured at the sampling locations and depths, that lay in the range of a mild hypoxia (25).

**Figure 1.**
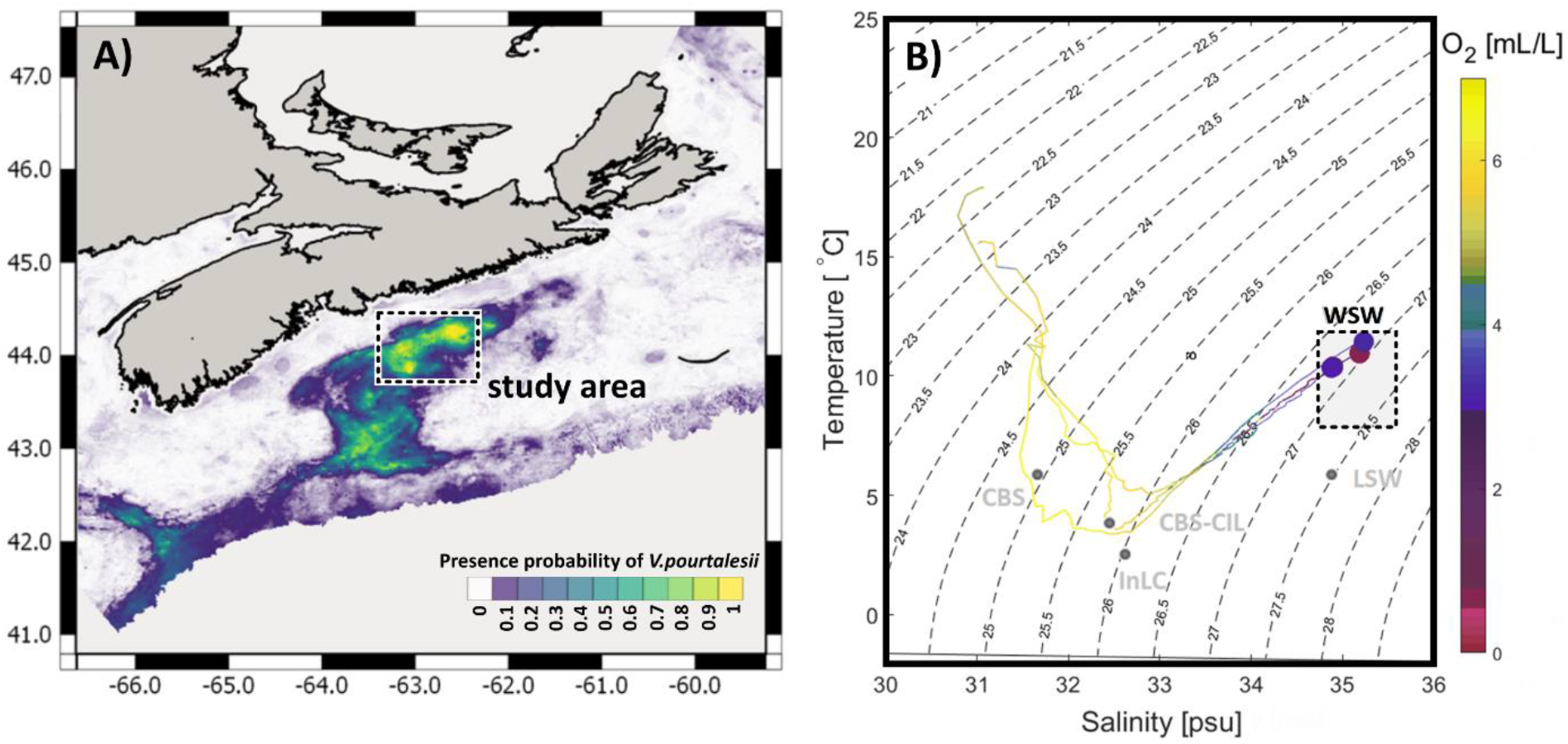
Map of sampling region on the Canadian shelf (A) and TS diagram (B). In (A) colours depict presence probability of *Vazella pourtalesii* based on data presented in Beazley et al. (17), with yellow indicating areas of highest occurrence probability. In B) colouring corresponds to oxygen concentrations measured during representative CTD casts at the study area. Water masses (light grey dots, labels and square) were added according to Dever et al. (101) and (102): CBS = Cabot Strait Subsurface Water; InLC= Inshore Labrador Current, CBS-CIL= Cold Intermediate Layer of Cabot Strait Subsurface Water, LSW= Labrador Slope Water, WSW=Warm Slope Water.

### Microscopic analyses of *V. pourtalesii*

In contrast to other sponges, Hexactinellidae mainly consist of a single syncytium, a fusion of eukaryotic cells forming multinucleate tissue, that permeates the whole sponge (12). By SEM we observed that the overall amount of sponge biomass in *V. pourtalesii* was low and its distribution within the spicule scaffolds was patchy (Figure 2A,B). Closer inspection of such biomass patches by light microscopy and by TEM microscopy (Figure 2C,D) revealed numerous host cells with their characteristic nuclei as well as high densities of microbial cells of various morphologies and with a dominance of comparatively small cell sizes (<1 μm). In addition, we frequently observed microbial cells that were attached to each other (Figure 2E,F).

**Figure 2.**
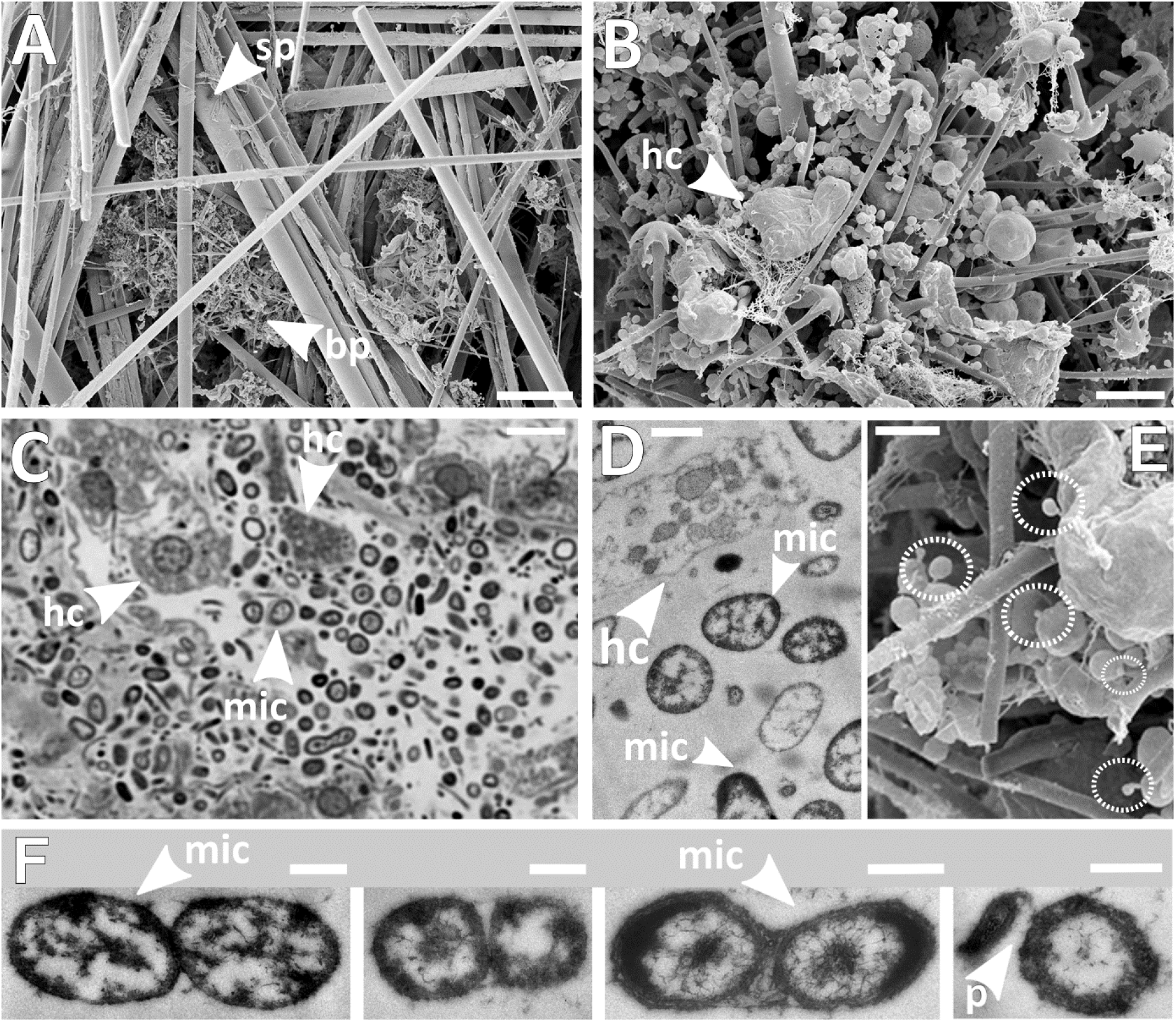
Microscopy of *Vazella pourtalesii* tissue. A) Scanning electron microscopy overview of spicule scaffolds (scale bar: 75 μm). B) SEM close-up image of a biomass patch (scale bar: 3 μm). C) Light-microscopy image (scale bar: 5 μm) and D) TEM image of the same biomass patch (scale bar: 1 μm). E) SEM close-up presumably showing smaller microbes attached to larger ones by stalk- or filament-like structures (scale bar: 1 μm). F) TEM microscopy images of adjacent microbial cells (scale bars: 500 nm). Acronyms: sp= spicule, bp = biomass patch, hc = host cell, mic= microbes, p = potential pilus.

### MAG selection

In total, 137 metagenome-assembled genomes (MAGs) of >50 % estimated completeness and <10 % redundancy were retrieved (Table S2). Proteobacteria followed by Patescibacteria were the dominant bacterial phyla according to amplicon analyses. In addition, LDA scores were obtained for MAGs based on their read abundance in the different metagenomic sample types (Figure S1): i) *V. pourtalesii* metagenomes vs. seawater metagenomes and ii) pristine *V. pourtalesii* vs. mooring *V. pourtalesii* vs. seawater metagenomes. Based on these assessments, we selected 13 representative *V. pourtalesii*-enriched MAGs for detailed analyses (Table 1). Five MAGs belonged to the candidate phylum Patescibacteria, three to the candidate phylum SAR324, four to the phylum Crenarchaeota, and one to the phylum Nanoarchaeota. The selected MAGs are evidently not redundant representations of the same microbial genomes, which is visualized by the comparatively long branches in the phylogenomic tree (Microscopy of *Vazella pourtalesii* tissue. A) Scanning electron microscopy overview of spicule scaffolds (scale bar: 75 μm). B) SEM close-up image of a biomass patch (scale bar: 3 μm). C) Light-microscopy image (scale bar: 5 μm) and D) TEM image of the same biomass patch (scale bar: 1 μm). E) SEM closeup presumably showing smaller microbes attached to larger ones by stalk- or filament-like structures (scale bar: 1 μm). F) TEM microscopy images of adjacent microbial cells (scale bars: 500 nm). Acronyms: sp= spicule, bp = biomass patch, hc = host cell, mic= microbes, p = potential pilus.

**Table 1.**
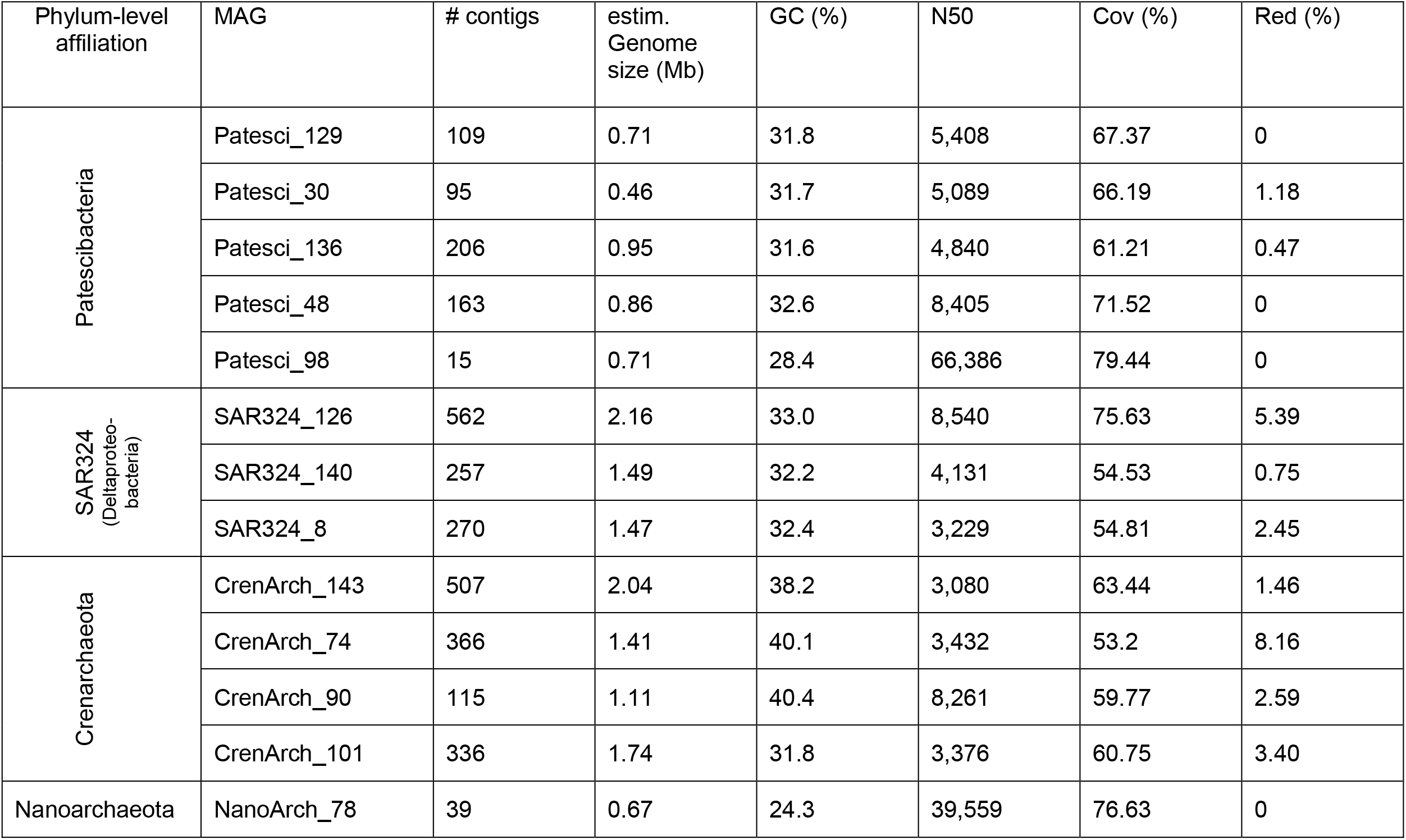
MAGs of the bacterial candidate phyla Patescibacteria and SAR324, and the archaeal phyla Crenarchaeota and Nanoarchaeota selected for detailed functional analysis. Genome properties were determined by QUAST, completeness and contamination estimations were performed by CheckM implemented in the metaWRAP pipeline. Acronyms: “Cov” – genome coverage, “Red” – redundancy.

Figure 3). Additionally, the symbiont MAGs showed a maximum of 86.8 %, 93.2 %, and 88.3 % similarity to each other in the ANI analysis for SAR324, Crenarchaeota and Patescibacteria, respectively (Table S4).

**Figure 3.**
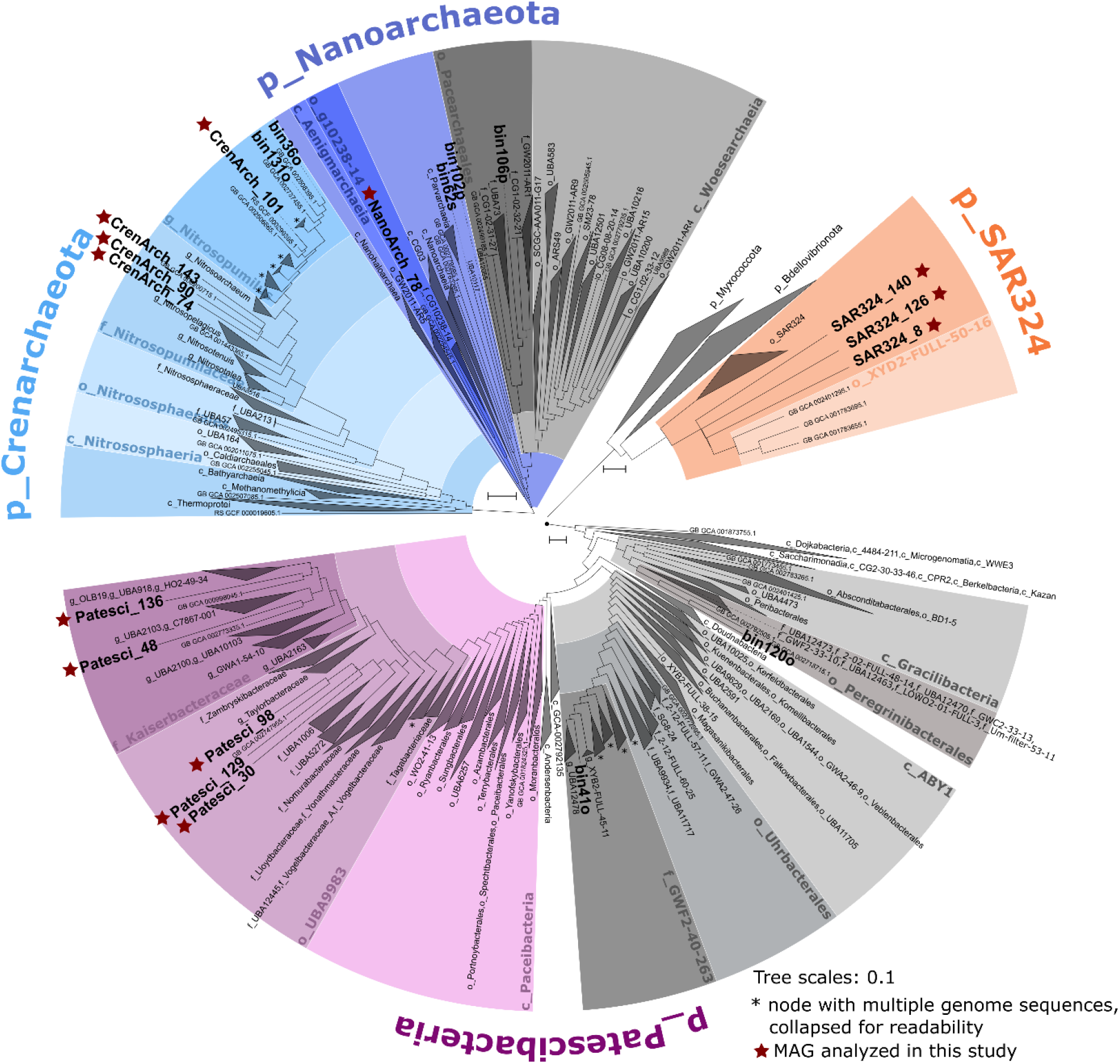
Subtrees of the GTDB-Tk phylogenetic tree showing the setting of the MAGs of this study (shown in bold) within the microbial phyla selected for detailed analysis. Class names are indicated by a leading “c_”, order names by “o_”, family names by “f_”, and genus names by “g_”. Vazella-enriched MAGs are marked with a red star.

### Phylogenetic placement of the major players

Three of the selected MAGs belong to the candidate phylum SAR324. The SAR324 clade was recently moved to the level of candidate phylum along with the publication of the whole genome-based classification of microbial genomes (26). The three SAR324 MAGs clustered together outside of known orders with relatively long branches, showing that they are genomically distinct from published genomes of their closest relatives (Microscopy of *Vazella pourtalesii* tissue. A) Scanning electron microscopy overview of spicule scaffolds (scale bar: 75 μm). B) SEM close-up image of a biomass patch (scale bar: 3 μm). C) Light-microscopy image (scale bar: 5 μm) and D) TEM image of the same biomass patch (scale bar: 1 μm). E) SEM close-up presumably showing smaller microbes attached to larger ones by stalk- or filament-like structures (scale bar: 1 μm). F) TEM microscopy images of adjacent microbial cells (scale bars: 500 nm). Acronyms: sp= spicule, bp = biomass patch, hc = host cell, mic= microbes, p = potential pilus.

Figure 3). In the amplicon analysis, this taxon was placed within the class Deltaproteobacteria, of which it posed the most abundant order (Figure S1 Linear discriminant analysis (LDA) Effect Size (LEfSe) plots of MAG abundance based on the abundance table calculated from read coverage data by the metaWRAP quant_bins module. Two sets of groups were analyzed: A) *Vazella* metagenomes vs. seawater reference metagenomes, and B) pristine *Vazella*-derived metagenomes (vazella_p) vs. mooring *Vazella*-derived metagenomes (vazella_m) vs. seawater reference metagenomes. An LDA score of 2 was selected as cut-off. Figure S2).

Three crenarchaeal MAGs clustered with *Cenarchaeum symbiosum* A (GCA000200715.1) associated with the sponge *Axinella mexicana* (27). One MAG (CrenArch_101) was closely related with *Nitrosopumilus* spp. isolated from Arctic marine sediment (28) and the deep-sea sponge *Neamphius huxleyi* (29). Crenarchaeota are not represented adequately in our amplicon data due to the sequencing primers bias towards bacteria.

Three of the five *V. pourtalesii*-enriched Patescibacteria MAGs clustered together with genome GCA002747955.1 from the oral metagenome of a dolphin (30). This clade is a sister group to other families of the order UBA9983 in the class Paceibacteria. The other two patescibacterial MAGs belonged to the family Kaiserbacteriaceae. They were placed separate from each other and outside of known genera, where they cluster with two different groundwater bacteria (GCA_000998045.1 and GCA_002773335.1). Patescibacteria were the second most abundant phylum in the amplicon data with a dominance of the class Parcubacteria which showed high abundances in *V. pourtalesii* compared to controls (Figure S3). The majority of the Parcubacteria (73%) remain unclassified, while 27% are classified as order Kaiserbacteria. As this phylum has not been noticed as particularily abundant in sponge microbiomes before, we tested whether they are sponge specific by comparison with the reference database of the Sponge Microbiome Project (SMP) (8), (data not shown). The 899 patescibacterial *V. pourtalesii* ASVs matched to 42 SMP OTUs (mostly listed as ‘unclassified Bacteria’ due to an older Silva version). We identified three SMP OTUs matching *V. pourtalesii* Kaiserbacteria ASVs, namely OTU0005080, OTU0007201, and OTU0159142. These OTUs occurred in 45, 26, and 7 sponge species, respectively, in the SMP reference database showing a global distribution.

The one Nanoarchaeota MAG NanoArch_78 belongs phylogenomically to the class Aenigmarchaeia, where it is placed together with MAG GCA_002254545.1 from a deep-sea hydrothermal vent sediment metagenome and outside of known families. Despite the primer bias towards bacteria in the amplicon analysis, the phylum Nanoarchaeota was among the most abundant microbial phyla in this analysis (Figure S1 Linear discriminant analysis (LDA) Effect Size (LEfSe) plots of MAG abundance based on the abundance table calculated from read coverage data by the metaWRAP quant_bins module. Two sets of groups were analyzed: A) *Vazella* metagenomes vs. seawater reference metagenomes, and B) pristine *Vazella*-derived metagenomes (vazella_p) vs. mooring *Vazella*-derived metagenomes (vazella_m) vs. seawater reference metagenomes. An LDA score of 2 was selected as cutoff. Figure S2).

### Genome sizes and GC contents

With respect to genome size, the MAGs range from 0.46 Mb (Patesci_30) to 2.16 Mb (SAR324_126), including completeness values into the genome size estimations, with N50 values between 3,080 (CrenArch_143) and 66,386 (Patesci_98) (Table 1). According to CheckM, between 53.2 % and 79.4 % of the genomes are covered and redundancies range from 0 % to 8.16 %. GC contents range from 24.3 % (NanoArch_78) to 40.38 % (CrenArch_90). We compared the MAGs to genomes of symbionts from other sponge species and of seawater-derived microbes of each respective phylum regarding their estimated genome sizes and GC contents (Figure 4, Table S3). Due to the lack of published genomes of SAR324 and Nanoarchaeota sponge symbionts, we included genome size and GC content data of unpublished symbionts of *Phakellia* spp. and *Stryphnus fortis* in this analysis. This comparison revealed that the genomes of *V. pourtalesii-enriched* microbes are exceptionally small with very low GC percentages. This trend is especially striking for Patescibacteria, SAR324 and Nanoarchaeota, as their values are not only low for sponge symbionts, but even below the levels of the respective related seawater microbes. While this is not the case for Crenarchaeota, the *V. pourtalesii* MAGs are, nevertheless, in the lower ranges regarding size and GC content in comparison to other sponge symbionts.

**Figure 4.**
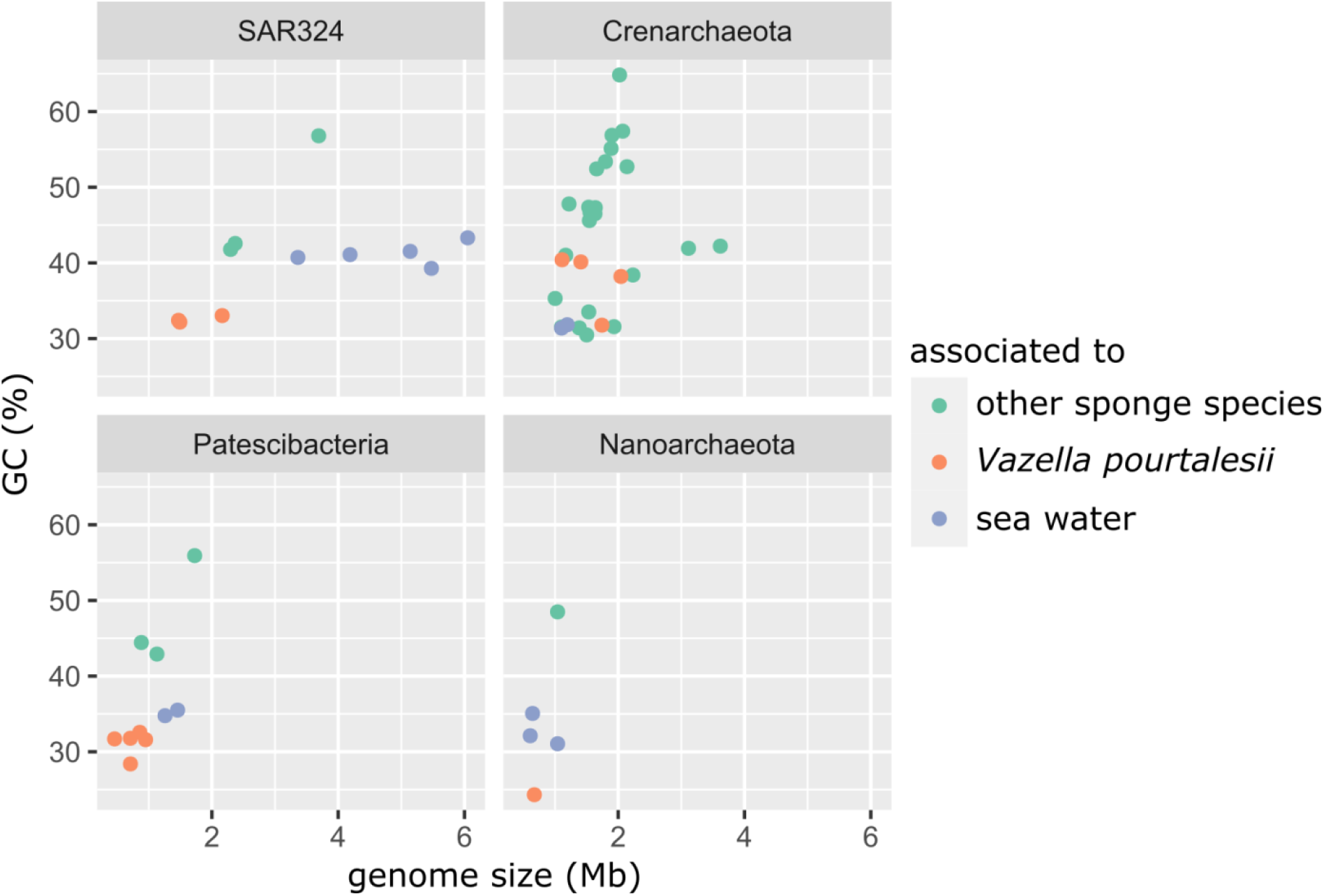
Comparison of the MAGs retrieved in this study to published MAGs from sponge and seawater metagenomes. From this study, only the MAGs enriched in either *Vazella pourtalesii* or in water were considered – ‘neutral’ ones were excluded from this analysis.

### Predicted lifestyle of the major players

#### SAR324

Metagenomic analysis of the three SAR324 MAGs from *V. pourtalesii* (Figure 5A) revealed the presence of a near-complete glycolysis pathway up to pyruvate (Pyr) along with the genes for the tricarboxylate acid (TCA) cycle and for conversions of the pentose phosphate pathway (PPP) (see Text S1 for in-depth analysis of more complex SAR324 and crenarchaeal MAGs). Pyruvate is converted aerobically by the pyruvate dehydrogenase enzyme complex into acetyl-CoA, which fuels the completely annotated (thiamin dependent) TCA cycle. While SAR324 have the genes for a near-complete respiratory chain, their lifestyle appears to be facultatively anaerobic. We detected enzymes of the glyoxylate-bypass (orange arrows within the TCA cycle in Figure 5A), which is required by bacteria to grow anaerobically on fatty acids and acetate (31). This is supported by the presence of a potential AMP-dependent acetyl-CoA synthetase to utilize acetate, whereas enzymes for fatty acids degradation were not found. SAR324 might gain additional energy by a cation-driven p-type ATPase and possibly also anaerobic respiration (furmarate, nitrite/sulfide respiration) (also see Text S1). There is evidence for assimilatory sulfate reduction, but the pathway was not fully resolved.

**Figure 5.**
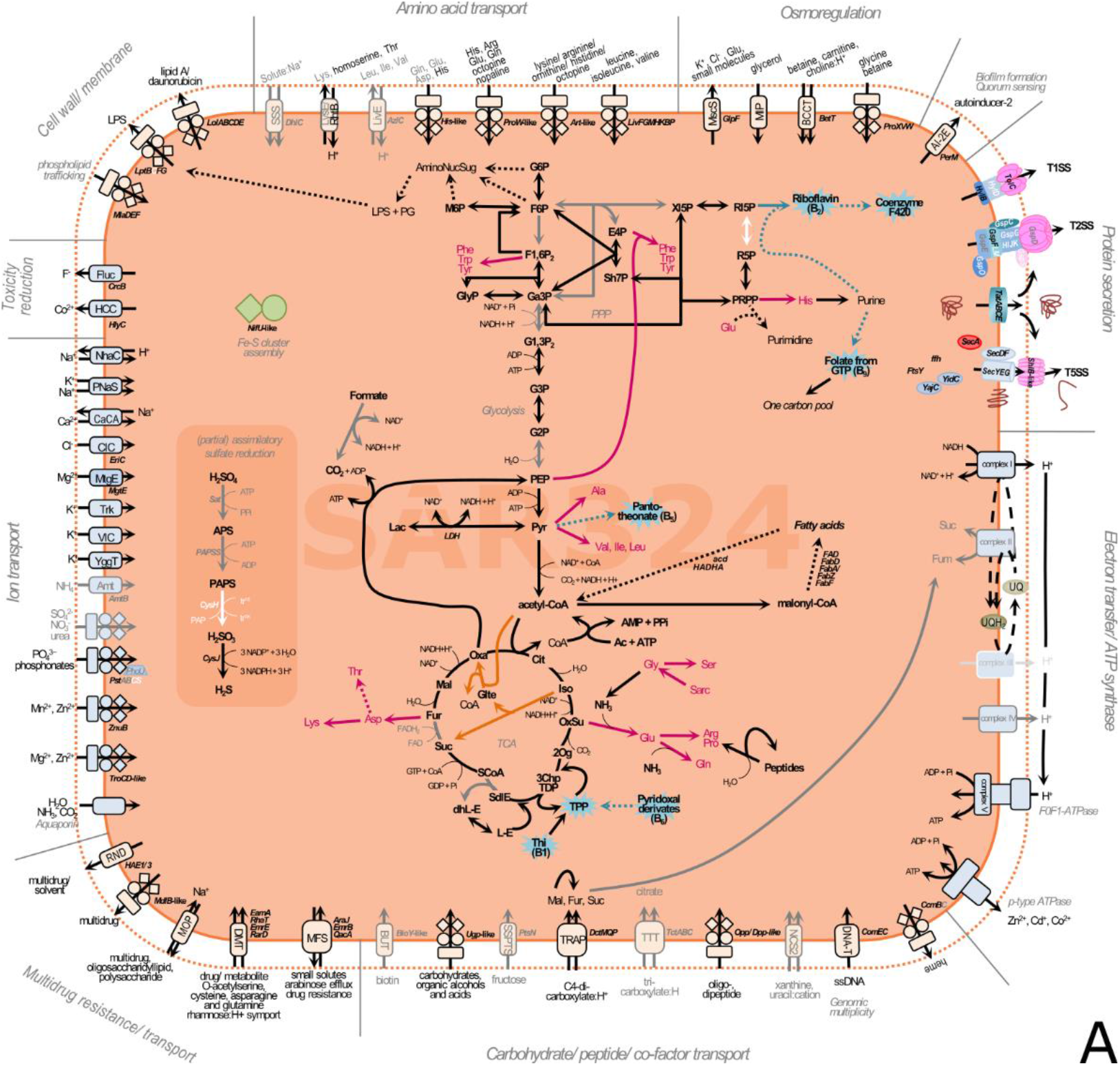
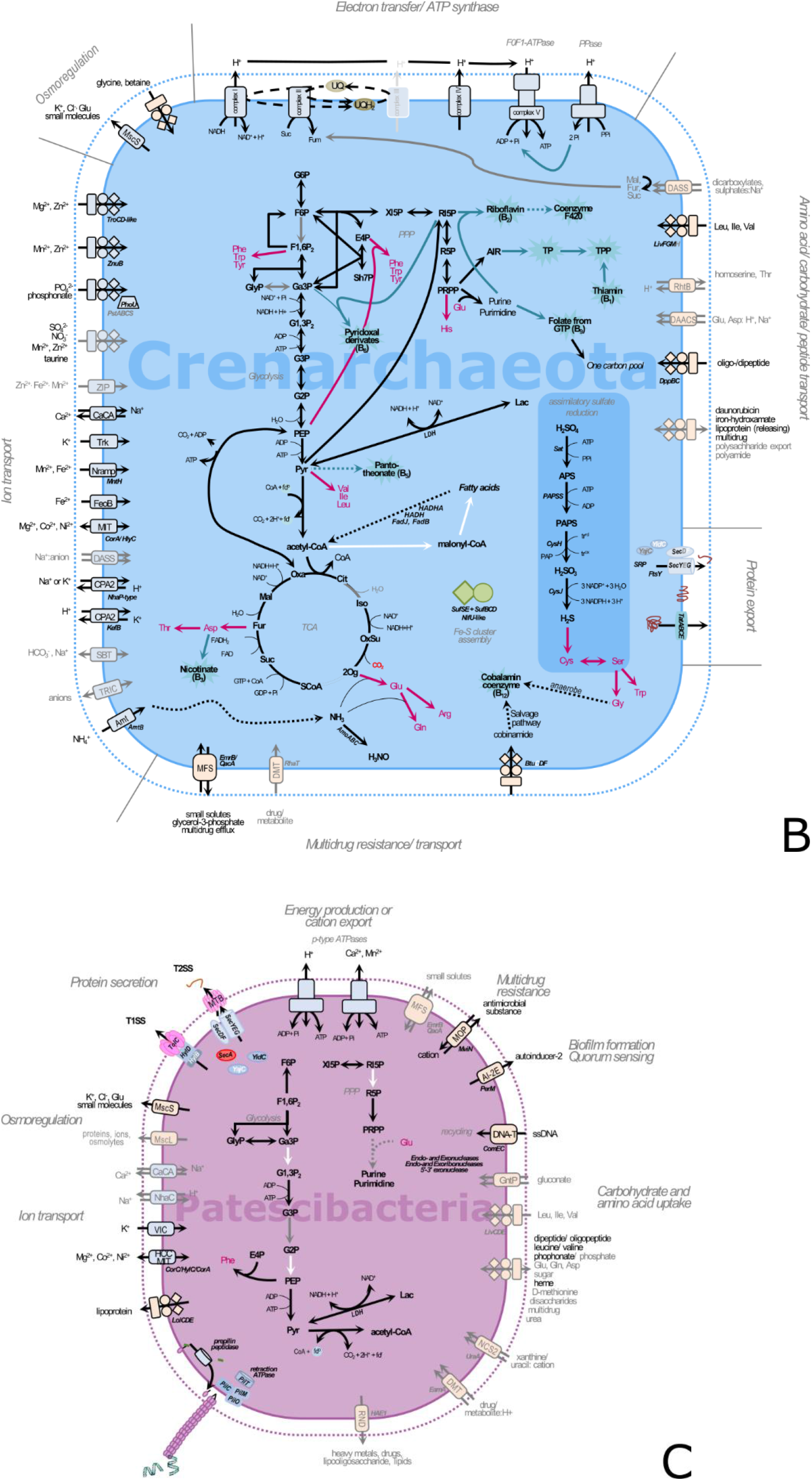
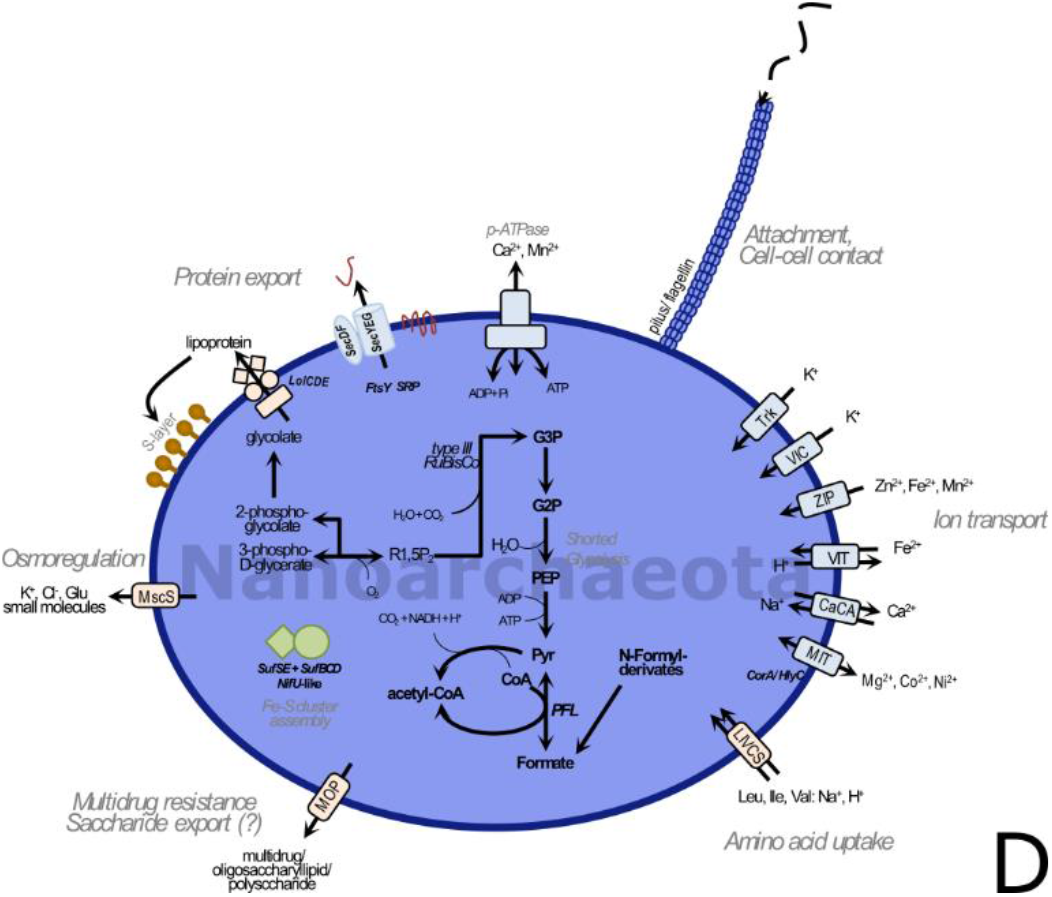
Reconstruction of metabolic features found in the genomes of A) SAR324, B) Crenarchaeota, C) Patescibacteria, and D) Nanoarchaeota. Solid lines indicate that genes/enzymes, or < 50% of a given pathway were found, dashed lines indicate less than 50% of a pathway were found. Grey arrows, writing and lining indicate that the genes/enzymes were found in less than 50% of genomes of the respective phylum. White arrows and writing indicate missing genes/enzymes. Co-factor synthesis is indicated by turquoise colour, amino acid production by magenta colour. Symport, antiport and direction are indicated by number and direction of arrows.

*V. pourtalesii*-associated SAR324 are able to take up di-and tricarboxylates using TRAP and TTT transporters, respectively. The imported substances can feed the TCA cycle under aerobic conditions or serve as energy source through fumarate respiration under anaerobic conditions (32, 33). The presence of lactate dehydrogenase (LHD) involved in fermentation is further supporting a facultatively anaerobic lifestyle (31).

SAR324 symbionts are capable of synthesizing diverse amino acids and b-vitamins (riboflavin, coenzyme F420, folate, panthotheonate and thiamin from pyridoxal), using precursors from glycolysis, PPP and TCA cycle (summarized in Figure 5A, see Text S1 for more details). Additionally, the genomes are well equipped with several transporters enabling the import and export of diverse substances (e.g., sugars, amino acids, petides, ions), (Text S1). These transporters are involved in, e.g. osmo-regulation and/or toxic ion reduction (cobalt, fluoride), and in multidrug resistance/import. A p-type ATPase was annotated that may aid in the export of cations or may use an electrochemical gradient for ATP synthesis. Proteins can be secreted by tat- and sec-transport as well as type 1 (T1SS) and type 2 (T2SS) secretion systems probably involved in excretion of symbiosis-relevant molecules. We further identified autoinducer-2 (AI-2) and DNA-T family transporter (Text S1).

#### Crenarchaeota

The Crenarchaeota of *V. pourtalesii* (Figure 5B) are capable of glycolysis from glucose-6-phosphate (G6P) to pyruvate (Pyr), which is likely anaerobically converted into acetyl-CoA using the enzyme pyruvate-ferredoxin oxidoreductase which suggests facultatively anaerobic metabolism. The TCA cycle was almost completely annotated. Genomic evidence for aerobic and anaerobic respiration (furmarate, nitrite/sulfide respiration) was detected (Text S1). Genes for autotrophic CO_2_ fixation in *V. pourtalesii*-associated symbionts were lacking, but assimilatory sulfate reduction was annotated completely. A PPase was annotated which might deliver phosphates to feed the ATP synthase) and in one MAG, a transporter to import dicarboxylates (fumarate, malate, succinate) was annotated, a feature we found in SAR324 as well. These substances could feed the TCA cycle under aerobic conditions. Additionally supporting the hypothesis of a facultatively anaerobic lifestyle is the presence of the enzyme lactate dehydrogenase (LHD) that is involved in fermentation (31). The crenarchaeal genomes encode the synthesis of an even greater number b-vitamins than the SAR324 genomes including riboflavin, coenzyme F420, folate, panthotheonate, pyridoxal, nicotinate and cobalmin (anaerobically) using precursors of central metabolism. Interestingly, they can synthesize thiamin from 5-phosphoribosyl diphosphate (PRPP), while SAR324 partially encode thiamin synthesis from pyridoxal which would need to be imported from an external source (e.g. from other community members).

The crenarchaeal genomes are further well equipped with transporters that facilitate import and export of diverse substances, such as sugars, amino acids, petides, and ions among others (Text S1). These transporters are involved in e.g. osmo-regulation, reflecting the adaptation to a saline environment, and in multidrug resistance/import. A p-type ATPase was annotated in the genomes that may be involved in the export of cations (forced by ATP utilization). Protein secretion can be realized by tat- and sec-transport, which might be involved in transport of proteins, such as those necessary for membrane formation and maintenance (Text S1).

#### Patescibacteria

The *V. pourtalesii-associated* Patescibacteria (Figure 5C) showed similar metabolic capacity to published Patescibacteria from other environments: While we found several enzymes involved in glycolysis, we could not resolve the pathway completely. The genomes lack enzymes involved in oxidative phosphorylation (respiration) and the TCA cycle. The synthesis pathways of the important precursor PRPP and subsequent synthesis of purines and pyrimidines were only partially encoded. The biosynthesis of phenylalanine (Phe) from phosphoenolpyruvate (PEP) and erythrose-4-phosphate (E4P) is encoded, but biosynthesis pathways for other amino acids, co-factors or vitamins are missing. We found two p-type ATPases which might export ions or provide energy (ATP) using cation and/or proton gradients present in the environment (holobiont). An anaerobic lifestyle is likely due to the presence of a lactate dehydrogenase (LDH), an enzyme involved in lactate fermentation and the anaerobic acetyl-CoA synthesis using the enzyme pyruvate-ferredoxin oxidoreductase. Patescibacterial MAGs also encode for some transporters, albeit in lower numbers compared to above-described SAR324 and crenarchaeal genomes. These transporters may be involved in osmo-regulation, multidrug in- and efflux, sugar-, amino acids-, and ion uptake. Regarding further symbiosis-relevant features, we detected the autoinducer transporter AI-2E. Additionally, we detected *ComEC/Rec2* and related proteins which are involved in the uptake of single stranded DNA (34). This is supported by the presence of *PilT,* the motor protein which is thought to drive pilin retraction prior to DNA uptake, and the pilus assembly proteins *PilM, PilO* and *PilC.* Even though not the full machinery was annotated, Patescibacteria may be able to take up foreign DNA via retraction of type IV-pilin-like structures into the periplasm and via *ComEC* through the inner membrane.

#### Nanoarchaeota

The nanoarchaeal MAG (Figure 5D) shows the genomic potential to convert glycerate-3-phosphate (G3P) to Pyr, which represents a shortened glycolysis pathway and results in reduced potential for energy production (ATP synthesis). It could use an anaerobic pyruvate-formate lyase (PFL) for acetyl-CoA production. Interestingly, an archaeal *type III* RuBisCo is encoded, which catalyzes light-independent CO_2_ fixation using ribulose-1,5-bisposphate and CO_2_ as substrates to synthesize G3P (35) to fill the only partially encoded glycolysis. All enzymes needed for respiration were absent supporting an anaerobic lifestyle. Energy might be gained using a cation-driven p-type ATPase. Like its published relatives (36, 37) the *V. pourtalesii*-associated nanorarchaeon lacks almost all known genes required for the *de novo* biosyntheses of amino acids, vitamins, nucleotides, cofactors, and lipids. The uptake of some amino acids and ions may be possible as few transporters were detected. Typical archaeal S-layer membrane proteins are encoded, which may be exported by an ABC-transporter (*LolCDE*) or by sec-transport. Other transporters are involved in osmo-regulation and in multidrug resistance and/or transport. We found a prepilin type IV leader peptidase encoded in the genome which is synthesized as a precursors before flagellin/pilin is incorporated into a filament (38).

## Discussion

### Microbial associations in *V. pourtalesii* syncytia

The glass sponge *V. pourtalesii* consists of a scaffold of spicules with cellular biomass concentrated in biomass patches that contain sponge as well as symbiont cells (Figure 2). We assume that the biomass patches were probably formed by dehydration of syncytial tissues during fixation resulting in higher biomass densities than in the *in vivo* situation (12). Surprisingly high numbers of microbial cells were found within the observed biomass patches, considering that microbes have rarely been noticed in glass sponges previously (12). In *V. pourtalesii,* the microbes appeared in various morphotypes indicating a taxonomically diverse microbial community. Microbial cells of strikingly small sizes (< 1μm) compared to those of shallow water demosponges (39) constituted a large fraction of the microbial community. Microbial cells were frequently seen in close association and even physically attached to each other (Figure 2E,F). These associations were observed between equally sized cells, but also between cells of distinctly different sizes, where the smaller microbes were attached to larger ones by stalk- or filament-like structures (Figure 2E).

### Main players in the *V. pourtalesii* microbial community with small, low GC genomes

While previously published sponge metagenomes and symbiont MAGs tended towards high GC contents (typically around 65-70%), the *V. pourtalesii* MAGs show lower GC levels in the range of 24-40% that are more similar to those of seawater metagenomes (40–42), (Figure 4, Table S3). Also genome sizes are on the smaller side in comparison to previously published sponge symbiont MAGs (40, 42). The large genome sizes of demosponge symbionts may be attributed to the specific genomic toolbox they require to utilize the mesohyl matrix, such as CAZy enzymes and arylsulfatases. These genes are frequently found enriched in the sponge symbiont genomes compared to free-living relatives (42, 43). This is, however, not the case in *V. pourtalesii* MAGs. On the contrary, here we see GC contents and genome sizes similar to and even below the ones of free-living marine microbes of the same respective phyla (Figure 4). Trophic specialization and avoidance of DNA replication cost have been proposed as hypotheses for genome reduction in free-living marine bacteria, e.g., of the genera *Idiomarina* (44) and *Pelagibacter* (45). For the *V. pourtalesii-associated* microbial community, the Black Queen hypothesis may best explain the apparent genomic streamlining: if some members carry out tasks that are beneficial to the whole microbial community, most other members will lose the ability to carry out these (often costly) tasks (46). The small sizes and low GC contents of the *V. pourtalesii* MAGs could, thus, be a sign of adaptation to generally nutrient-limiting environmental conditions (e.g. reviewed in (47)) and specialization on nutrient sources that are available within the sponge host environment, such as ammonia.

### The “givers” and “takers” hypothesis

Formerly placed within the Thaumarchaeota, the Crenarchaeota are well known and widespread sponge symbionts (48–51). Different genera have recently been observed in South Pacific Hexactinellida and Demospongiae (14). Also Nanoarchaeota and Patescibacteria have recently been noticed as members of sponge microbial communities including glass sponges, but no sponge-associated nanoarchaeal genome has been studied to date and – likely due to their low abundance in other sponge species – patescibacterial symbiont genomes have not been studied in detail (14, 52, 53). No sponge-derived genomes are available for the phylum SAR324 so far. The genomes of *V. pourtalesii* symbionts lack a number of properties that we know from typical shallow-water sponge symbionts (7), such as the potential for the production of secondary metabolites and arylsulfatases, and they are not enriched in genes encoding carbohydrate-active enzymes (CAZy). This underlines the above-stated hypothesis that these sponge symbionts do not need a diverse toolbox of genes to make use of a complex mesohyl like symbionts of Demospongiae. On the contrary, they seem to possess streamlined genomes to save resources and likely rely on each other for essential substances, such as certain amino acids and vitamins.

Based on the functional genetic content of the four symbiont phyla that we analyzed in greater detail, we propose two major strategies: “the givers”, namely SAR324 and Crenarchaeota and with comparatively larger, more complex genomes, and likely bigger in cell size, and “the takers”, Patescibacteria and Nanoarchaeota and with reduced genomes and likely smaller cell sizes. We posit that the givers – being genomically well equipped – could be producing and partly secreting all required amino acids and vitamins drawing energy from various aerobic as well as anaerobic processes (Figure 6). Regarding their metabolic repertoire, the here described Crenarchaeota are rather similar to the SAR324 bacteria, namely in their facultatively anaerobic lifestyle, the reactions of the central metabolism, and their ability for amino acid and vitamin biosynthesis. At the same time, while published sponge-associated or free-living Crenarchaeota have the genomic repertoire to fix carbon (e.g., (37, 50, 54–56)), such pathways appear to be absent in *V. pourtalesii*-associated Crenarchaeota. These findings indicate that the symbionts are specifically adapted to the conditions within their respective host sponge and the surrounding environment, e.g. low-oxygen conditions in this case.

**Figure 6.**
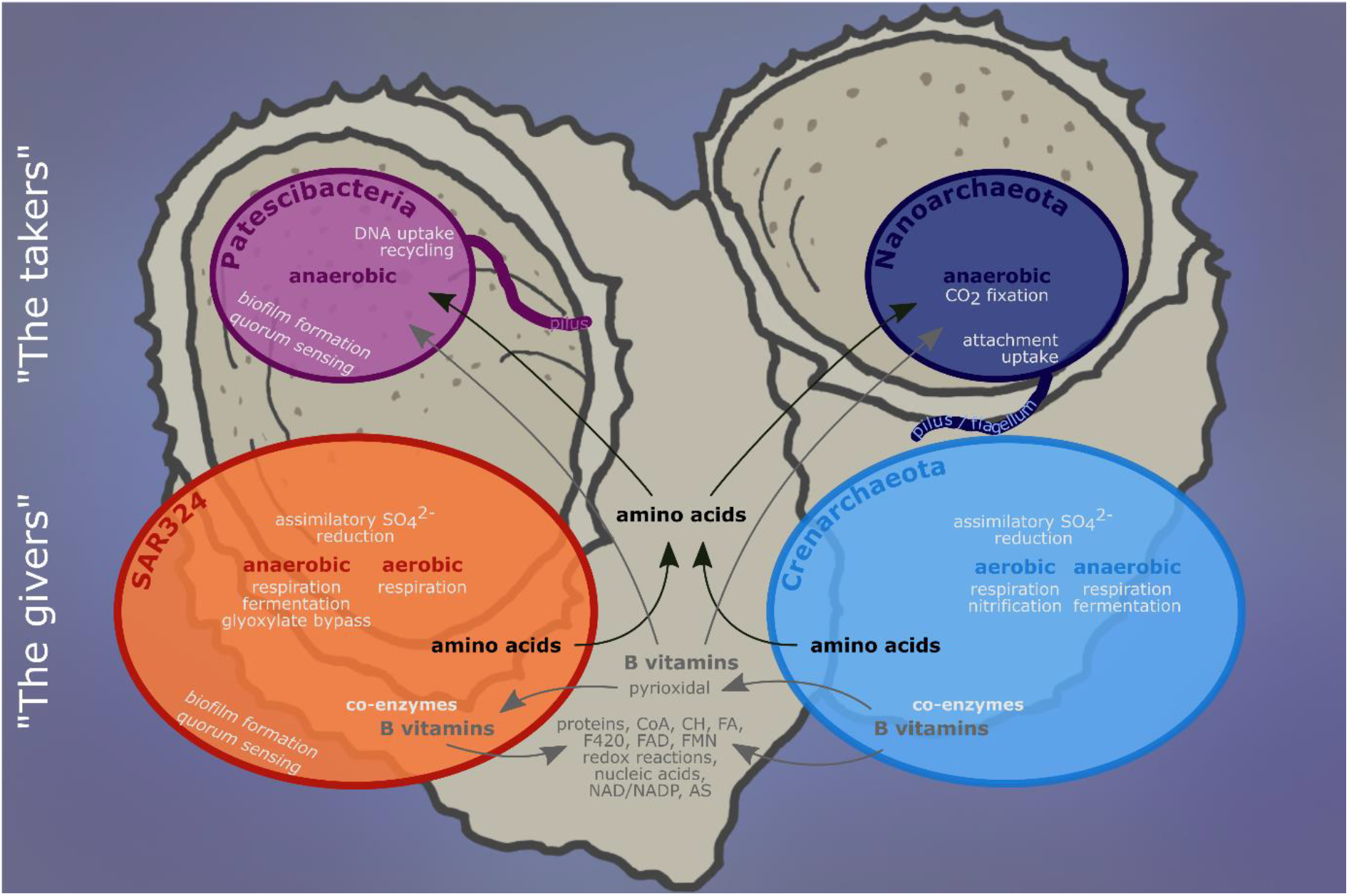
Summary model of the main metabolic interactions between the four microbial taxa studied herein. CoA: acetyl-CoA; CH: carbohydrates; FA: fatty acids; F420: coenzyme F420; AS: amino acids; FAD: flavin adenine dinucleotide; FMN: flavin mononucleotide; NAD: nicotinamide adenine dinucleotide; NADP: nicotinamide adenine dinucleotide phosphate.

Supporting our hypothesis of genome streamlining in the sense of the Black Queen hypothesis, the two “givers” also seem to depend on each other metabolically: Crenarchaeota can produce several b-vitamins, which might be used by members of SAR324. One example is pyrioxidal provided by Crenarchaeota to SAR324, which would thus be able to produce thiamin. Their genomic similarity, their difference to close relatives from other environments, and their metabolic interdependence reinforces our hypothesis that they are, in fact, symbionts specifically adapted to life within their *V. pourtalesii* host. Beyond the scope of the microbial community, microbial vitamin production may also have an important role in the animal host metabolism, such as the respiratory chain, the synthesis of coenzyme A, protein, fatty acid, nucleic acid and carbohydrate metabolism, and co-factor synthesis. As previously hypothesized for Demospongiae symbionts (7), the capacity for vitamin synthesis by microbes associated to *V. pourtalesii* might be an important factor in maintaining the symbiosis with the animal host.

It is tempting to speculate that the takers – reduced in size and functional potential – would scavange from their neighbors. Marine Patescibacteria are known (and named) for their reduced genomes and metabolic capacities (57–59) and also Nanoarchaeota are known for their dependence on a crenarchaeal host, although in very different marine environments such as hydrothermal vents (60, 61). Regarding the exchange of substances between microbes, we propose that the nanoarchaeal symbionts “ride” on the crenarchaeal symbionts, analogous to what is described for *Nanoarchaeum equitans* and *Ignicoccus hospitalis* (37, 62), which use pili for attachment and possibly also for metabolite uptake. Jahn et al. (62) showed that *Nanoarchaeum equitans* may get large amounts of lipids (and possibly other substances) from its associated *Ignicoccus hospitalis*. We hypothesize that the nanoarchaeum in *V. pourtalesii* might be likewise directly associated with the abundant Crenarchaeota and receive, e.g., lipids and DNA via cell attachment and using pili-like structures to maintain cell-cell contact (see, e.g., small cells attached to larger cells in Figure 2E), (37, 60, 63). Patescibacteria could take up required nutrients from their microenvironment, also utilizing pili equipped with a *pilT* motor protein and *comEC* enabling the uptake of DNA for the recycling of nucleotides (64) that they cannot build themselves. Interestingly, we detected copies of the *luxS* gene, the proposed autoinducer-2 (AI-2) synthetase, in patescibacterial and SAR324 genomes. There is strong evidence that Al-2E family homologues function as an AI-2 exporter in *E. coli* cells to control biofilm formation. AI-2 is a proposed signaling molecule for interspecies communication in bacteria (reviewed in (65)). Autoinducer production plays a crucial role in *Vibrio fischeri* colonization of (and maintenance in) the light organ of the host squid *Euprymna scolopes* (66) and was recently detected in sponge-associated *Vibrio* species (67). Microbial *quorom sensing* processes (such as biofilm formation, bioluminescence, motility, virulence factor secretion, antibiotic production, sporulation and competence for DNA uptake) (68) may display symbiosis-relevant features additional or alternative to the ones described before for sponge-associated microbes (e.g., arylsulfatases, TPRs, CRISPR-Cas).

## Conclusions

The present study aimed to characterize the diversity and function of microbes residing in the glass sponge *Vazella pourtalesii*. A general pattern emerged in that the *V. pourtalesii* symbionts displayed smaller genome sizes and lower GC contents than bacterial relatives from seawater or from demosponge symbionts. Genomic analysis revealed two putative functional strategies: the “givers” (SAR324 and Crenarchaeota) producing and most likely providing required amino acids and vitamins to the microbial community and the “takers” (Patescibacteria and Nanoarchaeota) depending on the provision of compounds like lipids and DNA that they likely take up via pili-like structures. Their localization within biomass patches together with the environmental low-oxygen conditions could serve as explanation for the unique compositional and functional properties of the microbial community of *V. pourtalesii*.

## Material and Methods

### Sampling and assessment of microbial community composition

Sampling was performed on a cruise to the Scotian Shelf off Nova Scotia, eastern Canadain August-September 2017 onboard CCGS *Martha L. Black (MLB2017001).* Here, we selected a subset of all samples received during this cruise (Table S1) to study the lifestyle strategies of the dominant members of the microbial community (for details on sampling, DNA extraction, and amplicon sequencing see (20)), where we cover the complete dataset to study the microbial diversity inside *V. pourtalesii* in response to anthropogenic activities). Briefly, sponge individuals were collected for this study from pristine areas by the remotely operated vehicle ROPOS (Canadian Scientific Submersible Facility, Victoria, Canada) and tissue subsamples were taken, rinsed in sterile filtered seawater and frozen at −80°C. Samples were collected at an average sampling depth of 168 m (min=161 m, max=183 m) which coincides with the base of the euphotic zone. Oceanographic data such as temperature, salinity and oxygen were collected using CTD casts (sensors by Sea-Bird Electronics SBE 25). Water samples were taken during CTD casts and using Niskin bottles of the ROPOS ROV. Additionally, samples were collected from an Ocean Tracking Network (OTN) acoustic mooring located approximately 10 km northwest of the Sambro Bank Sponge Conservation Area on the Scotian Shelf (20). The mooring was anchored ~ 5 m above the seabed and was deployed for ~ 13 months (15th of August 2016 – 5th of September 2017) prior to its recovery.

DNA was extracted using the DNeasy Power Soil Kit (Qiagen). After quantification and quality assessment by NanoDrop spectrophotometer and by PCR, the V3V4 variable regions of the 16S rRNA gene were amplified in a dual-barcoding approach (69) using a one-step PCR with the primers 5’-CCTACGGGAGGCAGCAG-3’ (70) and 5’-GGACTACHVGGGTWTCTAAT-3’ (71). Samples were sequenced on a MiSeq platform (MiSeqFGx, Ilumina) with v3 chemistry. The raw sequences were quality-filtered using BBDUK (BBMAP version 37.75 (72)) with a Q20, a minimum length of 250 nt. Sequences were processed in QIIME2 (versions 2018.6 and 2018.8 (73)) implementing the DADA2 algorithm (74) to determine Amplicon Sequence Variants (ASVs). Sequences were denoised and chimeras, chloroplast and mitochondrial sequences were removed. Taxonomy was assigned using a Naïve Bayes classifier (75) trained on the Silva 132 99% OTUs 16S database (76). An ASV-based phylogeny was generated using the FastTree2 plugin (77). The plots were produced with R (version 3.0.2 (78)), Inkscape (version 0.92.3 (79)), QGIS (version 2.18.4 (80)), and MATLAB (version R2016b including Gibbs Seawater toolbox (81)).

### Scanning Electron Microscopy

Tissue subsamples of three sponge individuals (Table S1) were fixed for scanning electron microscopy (SEM) onboard ship in 6.25% GDA in PBS (FisherScientific) in two technical replicates each. Samples were then washed 3x 15 min in PBS, post-fixed for 2 h in 2% osmiumtetroxide (Carl Roth, Germany) and washed again 3x 15 min in PBS. Samples were dehydrated in an ascending ethanol series (ROTIPURAN^®^ Carl Roth, Germany): 1×15 min 30% EtOH, 2×15 min 50% EtOH, 2×15 min 70% EtOH, 2×15 min 80% EtOH, 2×15 min 90% EtOH, 1×15 min 100% EtOH. Subsequent dehydration was continued with carbon dioxide in a Critical Point Dryer (BalzersCPD 030). After critical point drying, the samples were manually fractionated and sputter coated 3 min at 25 mA with gold/palladium (Balzers SCD 004). The preparations were visualized using a Hitachi S-4800 field emission scanning electron microscope (Hitachi High-Technologies Corporation, Tokyo, Japan) with a combination of upper and lower detector at an acceleration voltage of 3 kV and an emission current of 10 mA.

### Transmission Electron Microscopy and Light Microscopy

Tissue samples of three sponge individuals (Table S1) were fixed onboard ship in 2.5% glutaraldehyde in 0.1 M natriumcacodylate buffer (pH 7.4; Science Services GmbH) for transmission electron microscopy (TEM) and light microscopy in two technical replicates each. Samples were then rinsed 3x with buffer at 4°C, post-fixed for 2 h in 2% osmiumtetroxide (Carl Roth) and washed with buffer 3×15 min at 4°C. Samples were partially dehydrated with an ascending ethanol (ROTIPURAN^®^ Carl Roth) series (2×15 min 30% EtOH, 1×15 min 50% EtOH, up to 70% ethanol). Samples were stored at 4°C overnight before desilicification with 4% suprapure hydrofluoric acid (Merck; incubation of approximately 5 hours). Afterwards, samples were washed 8×15 min in 70% EtOH (with an overnight storage at 4°C in between washings). Dehydration was continued with a graded ethanol series (1×15 min 90% EtOH and 2×15 min 100% EtOH) followed by gradual infiltration with LR-White resin (Agar Scientific) at room temperature (1×1 h 2:1 Ethanol:LR-White; 1×1 h 1:1 Ethanol:LR-White; 1×1 h 1:2 Ethanol:LR-White; 2×2 h pure LR-White). Samples were incubated in pure LR-White resin at 4°C overnight before being transferred into fresh resin and polymerized in embedding capsules at 57°C for 2 days.

Semithin sections (0.5 μm) were cut (in technical replicates) sing an ultramicrotome (Reichert-Jung ULTRACUT E), equipped with a diamond knife (DIATOME, Switzerland), and afterwards stained with Richardson solution (ingredients from Carl Roth; prepared as described in (82)). Semithin sections were then mounted on SuperFrost Ultra Plus^®^ microscopy slices (Carl Roth; using Biomount medium produced by Plano) and visualized with an Axio Observer.Z1 microscope (Zeiss, Germany). Ultrathin sections (70 nm) were cut (in technical replicates) with the same ultramicrotome, mounted on pioloform coated copper grids (75 mesh; Plano) and contrasted with uranyl acetate (Science Services; 20 min incubation followed by washing steps with MilliQ water) and Reynold’s lead citrate (ingredients from Carl Roth; 3 min incubation followed by washing steps with MilliQ water). The ultrathin preparations were visualized at an acceleration voltage of 80 kV on a Tecnai G2 Spirit Bio Twin transmission electron microscope (FEI Company).

### Microbial functional repertoire

For metagenomic sequencing, DNA was extracted from the seven sponge samples (four from natural *Vazella* grounds and three from the mooring to optimize for differential coverage binning) and five seawater controls (Table S1) with the QiagenAllPrep DNA/RNA Mini Kit. Two washing steps with buffer AW2 were employed during DNA extraction. DNase and protease-free RNase A (Thermo Scientific) was used to remove remnant RNA from the DNA extracts. For seawater controls, DNA was extracted from one half of a PVDF membrane filter (seawater filter – SWF; see above). The DNA was concentrated by precipitation with 100 % ethanol and sodium acetate buffer and re-eluted in 50 μl water. For all extracts, DNA quantity and quality were assessed by NanoDrop measurements and Qubit assays, and 30 μl (diluted in water, if necessary) were sent for metagenomic Illumina Nextera sequencing (HiSeq 4000, 2×150 bp paired-end) at the Institute of Clinical Molecular Biology (IKMB) of Kiel University. Sequence quality of all read files was assessed with FastQC (83).

The raw reads were trimmed with Trimmomatic v0.36 (ILLUMINACLIP:NexteraPE-Pe.fa:2:30:10 LEADING:3 TRAILING:3 SLIDINGWINDOW:4:15 MINLEN:36) and coassembled with megahit v1.1.3 (84). The metaWRAP v1.0.2 pipeline was implemented for binning as follows (85): Initial binning was performed with metabat, metabat2, and maxbin2 (86–88) within metaWRAP. The bins were refined with the metaWRAP bin_refinement module and further improved where possible with the reassemble_bins module. This module uses the genome assembler SPAdes v3.12.0 (89) on two sets of reads mapped to the original bin with strict and more permissive settings and then compares the original bin with the two newly assembled genomes. Which of the three versions of the metagenome-assembled genome (MAG) was the best in each respective case and was, thus, used for further analyses, is indicated by the trailing letter in the names (‘o’ for ‘original’, ‘p’ for ‘permissive’, or ‘s’ for ‘strict’) in Table S2. The MAGs that were further analyzed in detail were renamed indicating their phylum-level affiliation and their bin number.

MAG taxonomy was determined by GTDB-Tk based on whole genome information and following the recently published, revised microbial taxonomy by Parks and colleagues (26, 90, 91). The phylogenomic trees produced by GTDB-Tk (77, 92, 93) were visualized on the Interactive Tree Of Life (iTOL) platform v4.3 (94). MAG abundance in the different metagenomic datasets was quantified with the metaWRAP quant_bins module and used to determine which MAGs were enriched in which sample type by calculating linear discriminant analysis (LDA) scores with LEfSe v1.0 (95) in two ways: i) “*V. pourtalesii”* vs. “water”, and ii) “pristine *V. pourtalesii”* vs. “mooring *V. pourtalesii”* vs. “water”. We identified MAGs belonging to the bacterial candidate phyla Patescibacteria and SAR324 and the archaeal phyla Crenarchaeota and Nanoarchaeota that were enriched in *V. pourtalesii* over seawater or in one of the *V. pourtalesii* subsets. The MAGs were compared to each other within their taxonomic groups using average nucleotide identity (ANI) of the pangenomic workflow of anvi’o v5.2 (96, 97) and they were compared to seawater and other host sponge-derived reference genomes (Table S3) regarding their genome sizes and GC contents. For functional annotations, interproscan v5.30-69.0 including GO term and pathway annotations was used (98, 99). The resulting EC numbers were converted to K terms to apply the online tool ‘Reconstruction Pathway’ in KEGG mapper (https://www.genome.jp/kegg/). Additionally, manual search in the annotation tables (DOI: 10.6084/m9.figshare.12280313) allowed the identification of several enzymes completing some pathways. Potential transporters were identified in the above-described annotation and additionally using the online tool TransportDB 2.0 (100).

## Data deposition

Detailed sample metadata was deposited in the PANGAEA database: https://doi.pangaea.de/10.1594/PANGAEA.917599. Amplicon and metagenomic raw read data were deposited in the NCBI database under BioProject PRJNA613976. Individual accession numbers for assembled MAGs are listed in Table S2. Interpro annotation output is available on figshare under DOI 10.6084/m9.figshare.12280313.

## Acknowledgments

The project “SponGES” has received funding from the European Union’s Horizon 2020 research and innovation programme under grant agreement No 679849. We thank our colleague Hans Tore Rapp for leading this great consortium for the last four years – you are dearly missed. Shiptime and Canadian participation was enabled by Fisheries and Oceans Canada’s (DFO) International Governance Strategy Science Program through project “Marine Biological Diversity Beyond Areas of National Jurisdiction (BBNJ): 3-Tiers of Diversity (Genes-Species-Communities)” led by EK (2017–2019). We appreciated the onboard support of the crew and scientific party of the expedition *MLB2017001*. Fred Whoriskey provided the OTN mooring samples. Andrea Hethke, Ina Clefsen, and the CRC1182 Z3 team (Katja Cloppenborg-Schmidt, Malte Rühlemann, John Baines) provided valuable support with the amplicon pipeline. Further, we want to thank the following people for microscopy related support: Yu-Chen Wu, Marie Sieberns, Anke Bleyer, Cay Kruse and Julia-Vanessa Böge.

## Supplementary material

Figure S1 Linear discriminant analysis (LDA) Effect Size (LEfSe) plots of MAG abundance based on the abundance table calculated from read coverage data by the metaWRAP quant_bins module. Two sets of groups were analyzed: A) *Vazella* metagenomes vs. seawater reference metagenomes, and B) pristine *Vazella*-derived metagenomes (vazella_p) vs. mooring *Vazella*-derived metagenomes (vazella_m) vs. seawater reference metagenomes. An LDA score of 2 was selected as cut-off.

Figure S2 The 15 most abundant microbial phyla (classes for Proteobacteria) in *V. pourtalesii,* representing > 99 % of the total microbiome. Microbial taxa are sorted after relative abundance in descending order from left to right. Symbols represent domain level taxonomic classification: circles indicate Bacteria, squares indicate Archaea. Patescibacteria (P), SAR324, and Nanoarchaeota (N) are particularly highlighted. The pie chart (grey) around the bubble chart indicates percentages of highest achievable taxonomic resolution for each taxonomically assignable Amplicon Sequence Variant (ASV). The asterisk highlights the three small pies next to it, which represent percentages of ASVs with highest achievable taxonomic resolution on phylum-level and kingdom-level, as well as sequences unassigned on phylum-level.

Figure S3 Heatmap of relative abundances [%] in our dataset of Patescibacteria classes within the phylum. Bubbles on the left side of the heatmap show richness (ASVs) of each class (max.=Parcubacteria 223 ASVs). Purple bars above the heatmap indicated relative abundance [%] of the phylum Patescibacteria within total microbiomes.

Table S1 Samples of this study and applied analyses.

Table S2 Overview of the MAGs binned from the *Vazella pourtalesii* metagenome in this study. Enrichment in one of the groups or all *Vazella* vs. water was determined by LefSe, classification is derived from GTDB-Tk, completeness and contamination were determined wich CheckM.

Table S3 Reference genomes for size and GC comparison.

Table S4 ANI analysis.

Text S1 Detailed description of metabolic features detected in SAR324 and Crenarchaeota.

## References

1. Maldonado M, Aguilar R, Bannister RJ, Bell JJ, Conway KW, Dayton PK, Díaz C, Gutt J, Kelly M, Kenchington ELR, Leys SP, Pomponi SA, Rapp HT, Rützler K, Tendal OS, Vacelet J, Young CM. 2015. Sponge grounds as key marine habitats: A synthetic review of types, structure, functional roles, and conservation concerns, p. 1–39. In Rossi, S, Bramanti, L, Gori, A, Orejas, C (eds.), Marine Animal Forests. Springer International Publishing, Cham.

2. Howell K-L, Piechaud N, Downie A-L, Kenny A. 2016. The distribution of deep-sea sponge aggregations in the North Atlantic and implications for their effective spatial management. Deep Sea Res Part I Oceanogr Res Pap 115:309–320.

3. Beazley L, Kenchington E, Yashayaev I, Murillo FJ. 2015. Drivers of epibenthic megafaunal composition in the sponge grounds of the Sackville Spur, northwest Atlantic. Deep Res Part I Oceanogr Res Pap 98:102–114.

4. Hawkes N, Korabik M, Beazley L, Rapp HT, Xavier JR, Kenchington E. 2019. Glass sponge grounds on the Scotian Shelf and their associated biodiversity. Mar Ecol Prog Ser 614:91–109.

5. Murillo FJ, Kenchington E, Koen-Alonso M, Guijarro J, Kenchington TJ, Sacau M, Beazley L, Rapp HT. 2020. Mapping benthic ecological diversity and interactions with bottom-contact fishing on the Flemish Cap (northwest Atlantic). Ecol Indic 112:106135.

6. Thomas T, Moitinho-Silva L, Lurgi M, Björk JR, Easson C, Astudillo-García C, Olson JB, Erwin PM, Lóspez-Legentil S, Luter H, Chaves-Fonnegra A, Costa R, Schupp PJ, Steindler L, Erpenbeck D, Gilbert J, Knight R, Ackermann G, Lopez JV, Taylor MW, Thacker RW, Montoya JM, Hentschel U, Webster NS. 2016. Diversity, structure and convergent evolution of the global sponge microbiome. Nat Commun 7:11870.

7. Pita L, Rix L, Slaby BM, Franke A, Hentschel U. 2018. The sponge holobiont in a changing ocean: from microbes to ecosystems. Microbiome 6:46.

8. Moitinho-Silva L, Nielsen S, Amir A, Gonzalez A, Ackermann GL, Cerrano C, Astudillo-García C, Easson C, Sipkema D, Liu F, Steinert G, Kotoulas G, McCormack GP, Feng G, Bell JJ, Vicente J, Björk JR, Montoya JM, Olson JB, Reveillaud J, Steindler L, Pineda M-C, Marra MV, Ilan M, Taylor MW, Polymenakou P, Erwin PM, Schupp PJ, Simister RL, Knight R, Thacker RW, Costa R, Hill RT, Lopez-Legentil S, Dailianis T, Ravasi T, Hentschel U, Li Z, Webster NS, Thomas T. 2017. The sponge microbiome project. Gigascience 6:gix077.

9. Maldonado M, Ribes M, van Duyl FC. 2012. Nutrient fluxes through sponges: Biology, budgets, and ecological implications, p. 113–182. In Becerro M, Uriz M, Maldonado M, Turon X (eds.), Advances in Marine Biology, Volume 62. Elsevier Ltd. Academic Press, Amsterdam.

10. de Goeij JM, van Oevelen D, Vermeij MJA, Osinga R, Middelburg JJ, de Goeij AFPM, Admiraal W. 2013. Surviving in a marine desert: The sponge loop retains resources within coral reefs. Science 342:108–110.

11. Krautter M, Conway KW, Barrie JV, Neuweiler M. 2001. Discovery of a “Living Dinosaur”: Globally unique modern hexactinellid sponge reefs off British Columbia, Canada. Facies 44:265–282.

12. Leys SP, Mackie GO, Reiswig HM. 2007. The Biology of Glass Sponges, p. 1–145. In Sims D (ed.), Advances in Marine Biology, Volume 52. Elsevier Ltd. Academic Press.

13. van Soest RWM, Boury-Esnault N, Vacelet J, Dohrmann M, Erpenbeck D, de Voogd NJ, Santodomingo N, Vanhoorne B, Kelly M, Hooper JNA. 2012. Global diversity of sponges (Porifera). PLoS One 7:e35105.

14. Steinert G, Busch K, Bayer K, Kodami S, Arbizu PM, Kelly M, Mills S, Erpenbeck D, Dohrmann M, Wörheide G, Hentschel U, Schupp PJ. 2020. Compositional and Quantitative Insights Into Bacterial and Archaeal Communities of South Pacific DeepSea Sponges (Demospongiae and Hexactinellida). Front Microbiol 11:716.

15. Tian R-M, Sun J, Cai L, Zhang W-P, Zhou G-W, Qiu J-W, Qian P-Y. 2016. The deepsea glass sponge *Lophophysema eversa* harbours potential symbionts responsible for the nutrient conversions of carbon, nitrogen and sulfur. Environ Microbiol 18:2481–2494.

16. Schmidt O. 1870. Grundzüge einer Spongien-Fauna des atlantischen Gebietes. Wilhelm Engelmann, Leipzig.

17. Beazley L, Wang Z, Kenchington E, Yashayaev I, Rapp HT, Xavier JR, Murillo FJ, Fenton D, Fuller S. 2018. Predicted distribution of the glass sponge *Vazella pourtalesi* on the Scotian Shelf and its persistence in the face of climatic variability. PLoS One 13:e0205505.

18. Townsend DW, Pettigrew NR, Thomas MA, Neary MG, McGillicuddy Jr. DJ, O’Donnell J. 2015. Water masses and nutrient sources to the Gulf of Maine. J Mar Res 73:93–122.

19. Hendry KR, Cassarino L, Bates SL, Culwick T, Frost M, Goodwin C, Howell KL. 2019. Silicon isotopic systematics of deep-sea sponge grounds in the North Atlantic. Quat Sci Rev 210:1–14.

20. Busch K, Beazley L, Kenchington E, Whoriskey F, Slaby B, Hentschel U. Microbial diversity of the glass sponge *Vazella pourtalesii* in response to anthropogenic activities. In review at Conservation Genetics. bioRxiv preprint DOI 10.1101/2020.05.19.102806.

21. Dohrmann M, Wörheide G. 2017. Dating early animal evolution using phylogenomic data. Sci Rep 7:3599.

22. Mentel M, Röttger M, Leys S, Tielens AGM, Martin WF. 2014. Of early animals, anaerobic mitochondria, and a modern sponge. BioEssays 36:924–932.

23. Mills DB, Francis WR, Vargas S, Larsen M, Elemans CP, Canfield DE, Wörheide G. 2018. The last common ancestor of animals lacked the HIF pathway and respired in low-oxygen environments. eLife 7:e31176.

24. Leys SP, Kahn AS. 2018. Oxygen and the Energetic Requirements of the First Multicellular Animals. Integr Comp Biol 58:666–676.

25. Hofmann AF, Peltzer ET, Walz PM, Brewer PG. 2011. Hypoxia by degrees: Establishing definitions for a changing ocean. Deep Sea Res Part I Oceanogr Res Pap 58:1212–1226.

26. Parks DH, Chuvochina M, Waite DW, Rinke C, Skarshewski A, Chaumeil P-A, Hugenholtz P. 2018. A standardized bacterial taxonomy based on genome phylogeny substantially revises the tree of life. Nat Biotechnol 36:996–1004.

27. Hallam SJ, Konstantinidis KT, Putnam N, Schleper C, Watanabe Y, Sugahara J, Preston C, de la Torre J, Richardson PM, DeLong EF. 2006. Genomic analysis of the uncultivated marine crenarchaeote *Cenarchaeum symbiosum*. Proc Natl Acad Sci U S A 103:18296–18301.

28. Park S-J, Kim J-G, Jung M-Y, Kim S-J, Cha I-T, Ghai R, Martín-Cuadrado A-B, Rodríguez-Valera F, Rhee S-K. 2012. Draft Genome Sequence of an Ammonia-Oxidizing Archaeon, “*Candidatus* Nitrosopumilus sediminis” AR2, from Svalbard in the Arctic Circle. J Bacteriol 194:6948–6949.

29. Parks DH, Rinke C, Chuvochina M, Chaumeil P-A, Woodcroft BJ, Evans PN, Hugenholtz P, Tyson GW. 2017. Recovery of nearly 8,000 metagenome-assembled genomes substantially expands the tree of life. Nat Microbiol 2:1533–1542.

30. Dudek NK, Sun CL, Burstein D, Kantor RS, Aliaga Goltsman DS, Bik EM, Thomas BC, Banfield JF, Relman DA. 2017. Novel Microbial Diversity and Functional Potential in the Marine Mammal Oral Microbiome. Curr Biol 27:3752–3762.

31. White D, Drummond J, Fuqua C. 2012. The Physiology and Biochemistry of Prokaryotes. Oxford University Press.

32. Unden G, Strecker A, Kleefeld A, Kim O Bin. 2016. C4-Dicarboxylate Utilization in Aerobic and Anaerobic Growth. Ecosal Plus 7:1–33.

33. Rosa LT, Bianconi ME, Thomas GH, Kelly DJ. 2018. Tripartite ATP-Independent Periplasmic (TRAP) Transporters and Tripartite Tricarboxylate Transporters (TTT): From Uptake to Pathogenicity. Front Cell Infect Microbiol 8:33.

34. Draskovic I, Dubnau D. 2005. Biogenesis of a putative channel protein, ComEC, required for DNA uptake: membrane topology, oligomerization and formation of disulphide bonds. Mol Microbiol 55:881–896.

35. Sato T, Atomi H, Imanaka T. 2007. Archaeal type III RuBisCOs function in a pathway for AMP metabolism. Science 315:1003–1006.

36. Waters E, Hohn MJ, Ahel I, Graham DE, Adams MD, Barnstead M, Beeson KY, Bibbs L, Bolanos R, Keller M, Kretz K, Lin X, Mathur E, Ni J, Podar M, Richardson T, Sutton GG, Simon M, Söll D, Stetter KO, Short JM, Noordewier M. 2003. The genome of *Nanoarchaeum equitans:* Insights into early archaeal evolution and derived parasitism. Proc Natl Acad Sci 100:12984–12988.

37. Podar M, Anderson I, Makarova KS, Elkins JG, Ivanova N, Wall MA, Lykidis A, Mavromatis K, Sun H, Hudson ME, Chen W, Deciu C, Hutchison D, Eads JR, Anderson A, Fernandes F, Szeto E, Lapidus A, Kyrpides NC, Saier MH, Richardson PM, Rachel R, Huber H, Eisen JA, Koonin EV, Keller M, Stetter KO. 2008. A genomic analysis of the archaeal system *Ignicoccus hospitalis*-*Nanoarchaeum equitans*. Genome Biol 9:R158.

38. Bardy SL, Jarrell KF. 2003. Cleavage of preflagellins by an aspartic acid signal peptidase is essential for flagellation in the archaeon *Methanococcus voltae*. Mol Microbiol 50:1339–1347.

39. Gloeckner V, Wehrl M, Moitinho-Silva L, Gernert C, Schupp P, Pawlik JR, Lindquist NL, Erpenbeck D, Wörheide G, Hentschel U. 2014. The HMA-LMA dichotomy revisited: An electron microscopical survey of 56 sponge species. Biol Bull 227:78–88.

40. Moreno-Pino M, Cristi A, Gillooly JF, Trefault N. 2020. Characterizing the microbiomes of Antarctic sponges: a functional metagenomic approach. Sci Rep 10:645.

41. Horn H, Slaby BM, Jahn MT, Bayer K, Moitinho-Silva L, Förster F, Abdelmohsen UR, Hentschel U. 2016. An enrichment of CRISPR and other defense-related features in marine sponge-associated microbial metagenomes. Front Microbiol 7:1751.

42. Slaby BM, Hackl T, Horn H, Bayer K, Hentschel U. 2017. Metagenomic binning of a marine sponge microbiome reveals unity in defense but metabolic specialization. ISME J 11:2465–2478.

43. Kamke J, Sczyrba A, Ivanova N, Schwientek P, Rinke C, Mavromatis K, Woyke T, Hentschel U. 2013. Single-cell genomics reveals complex carbohydrate degradation patterns in poribacterial symbionts of marine sponges. ISME J 7:2287–2300.

44. Qin Q-L, Li Y, Sun L-L, Wang Z-B, Wang S, Chen X-L, Oren A, Zhang Y-Z. 2019. Trophic Specialization Results in Genomic Reduction in Free-Living Marine *Idiomarina* Bacteria. mBio 10:e02545–18.

45. Giovannoni SJ, Tripp HJ, Givan S, Podar M, Vergin KL, Baptista D, Bibbs L, Eads J, Richardson TH, Noordewier M, Rappé MS, Short JM, Carrington JC, Mathur EJ. 2005. Genome Streamlining in a Cosmopolitan Oceanic Bacterium. Science 309:1242–1245.

46. Morris JJ, Lenski RE, Zinser ER. 2012. The Black Queen Hypothesis: Evolution of Dependencies through Adaptive Gene Loss. mBio 3:e00036–12.

47. Dutta C, Paul S. 2012. Microbial lifestyle and genome signatures. Curr Genomics 13:153–162.

48. Bayer K, Schmitt S, Hentschel U. 2008. Physiology, phylogeny and *in situ* evidence for bacterial and archaeal nitrifiers in the marine sponge *Aplysina aerophoba*. Environ Microbiol 10:2942–2955.

49. Moeller FU, Webster NS, Herbold CW, Behnam F, Domman D, Albertsen M, Mooshammer M, Markert S, Turaev D, Becher D, Rattei T, Schweder T, Richter A, Watzka M, Nielsen PH, Wagner M. 2019. Characterization of a thaumarchaeal symbiont that drives incomplete nitrification in the tropical sponge *Ianthella basta*. Environ Microbiol 21:3831–3854.

50. Moitinho-Silva L, Díez-Vives C, Batani G, Esteves AIS, Jahn MT, Thomas T. 2017. Integrated metabolism in sponge–microbe symbiosis revealed by genome-centered metatranscriptomics. ISME J 11:1–16.

51. Zhang S, Song W, Wemheuer B, Reveillaud J, Webster N, Thomas T. 2019. Comparative Genomics Reveals Ecological and Evolutionary Insights into Sponge-Associated Thaumarchaeota. mSystems 4:e00288–19.

52. Engelberts JP, Robbins SJ, de Goeij JM, Aranda M, Bell SC, Webster NS. 2020. Characterization of a sponge microbiome using an integrative genome-centric approach. ISME J 14:1100–1110.

53. Turon M, Uriz MJ. 2020. New Insights Into the Archaeal Consortium of Tropical Sponges. Front Mar Sci 6:789.

54. Berg IA. 2011. Ecological aspects of the distribution of different autotrophic CO2 fixation pathways. Appl Environ Microbiol 77:1925–36.

55. Bayer B, Vojvoda J, Offre P, Alves RJE, Elisabeth NH, Garcia JAL, Volland J-M, Srivastava A, Schleper C, Herndl GJ. 2016. Physiological and genomic characterization of two novel marine thaumarchaeal strains indicates niche differentiation. ISME J 10:1051–1063.

56. Hügler M, Sievert SM. 2011. Beyond the Calvin Cycle: Autotrophic Carbon Fixation in the Ocean. Ann Rev Mar Sci 3:261–289.

57. Rinke C, Schwientek P, Sczyrba A, Ivanova NN, Anderson IJ, Cheng J-F, Darling AE, Malfatti S, Swan BK, Gies EA, Dodsworth JA, Hedlund BP, Tsiamis G, Sievert SM, Liu W-T, Eisen JA, Hallam SJ, Kyrpides NC, Stepanauskas R, Rubin EM, Hugenholtz P, Woyke T. 2013. Insights into the phylogeny and coding potential of microbial dark matter. Nature 499:431–437.

58. Woyke T, Doud DFR, Eloe-Fadrosh EA. 2019. Genomes From Uncultivated Microorganisms, p. 437–442. In Encyclopedia of Microbiology. Elsevier.

59. Castelle CJ, Brown CT, Anantharaman K, Probst AJ, Huang RH, Banfield JF. 2018. Biosynthetic capacity, metabolic variety and unusual biology in the CPR and DPANN radiations. Nat Rev Microbiol 16:629–645.

60. Forterre P, Gribaldo S, Brochier-Armanet C. 2009. Happy together: genomic insights into the unique *Nanoarchaeum*/*Ignicoccus* association. J Biol 8:7.

61. Nicks T, Rahn-Lee L. 2017. Inside Out: Archaeal Ectosymbionts Suggest a Second Model of Reduced-Genome Evolution. Front Microbiol 8:384.

62. Jahn U, Summons R, Sturt H, Grosjean E, Huber H. 2004. Composition of the lipids of *Nanoarchaeum equitans* and their origin from its host *Ignicoccus* sp. strain KIN4/I. Arch Microbiol 182:404–413.

63. Albers S-V, Meyer BH. 2011. The archaeal cell envelope. Nat Rev Microbiol 9:414–426.

64. Mell JC, Redfield RJ. 2014. Natural Competence and the Evolution of DNA Uptake Specificity. J Bacteriol 196:1471 –1483.

65. Miller MB, Bassler BL. 2001. Quorum Sensing in Bacteria. Annu Rev Microbiol 55:165–199.

66. Chun CK, Troll JV, Koroleva I, Brown B, Manzella L, Snir E, Almabrazi H, Scheetz TE, de Fatima Bonaldo M, Casavant TL, Soares MB, Ruby EG, McFall-Ngai MJ. 2008. Effects of Colonization, Luminescence, and Autoinducer on Host Transcription During Development of the Squid-*Vibrio* Association. Proc Natl Acad Sci 105:11323–11328.

67. Zan J, Fuqua C, Hill RT. 2011. Diversity and functional analysis of *luxS* genes in Vibrios from marine sponges *Mycale laxissima* and *Ircinia strobilina*. ISME J 5:1505–1516.

68. Ng W-L, Bassler BL. 2009. Bacterial Quorum-Sensing Network Architectures. Annu Rev Genet 43:197–222.

69. Kozich JJ, Westcott SL, Baxter NT, Highlander SK, Schloss PD. 2013. Development of a dual-index sequencing strategy and curation pipeline for analyzing amplicon sequence data on the miseq illumina sequencing platform. Appl Environ Microbiol 79:5112–5120.

70. Muyzer G, de Waal EC, Uitterlinden AG. 1993. Profiling of complex microbial populations by denaturing gradient gel electrophoresis analysis of polymerase chain reaction-amplified genes coding for 16S rRNA. Appl Environ Microbiol 59:695–700.

71. Caporaso JG, Lauber CL, Walters WA, Berg-Lyons D, Lozupone CA, Turnbaugh PJ, Fierer N, Knight R. 2011. Global patterns of 16S rRNA diversity at a depth of millions of sequences per sample. Proc Natl Acad Sci 108:4516–4522.

72. Bushnell B. 2017. BBMap short read aligner, and other bioinformatic tools. Available online at: https://sourceforge.net/projects/bbmap/.

73. Bolyen E, Rideout JR, Dillon MR, Bokulich NA, Abnet CC, Al-Ghalith GA, Alexander H, Alm EJ, Arumugam M, Asnicar F, Bai Y, Bisanz JE, Bittinger K, Brejnrod A, Brislawn, CJ, Brown CT, Callahan BJ, Caraballo-Rodríguez AM, Chase J, Cope EK, DaSilva R, Diener C, Dorrestein PC, Douglas GM, Durall DM, Duvallet C, Edwardson CF, Ernst M, Estaki M, Fouquier J, Gauglitz JM, Gibbons SM, Gibson DL, Gonzalez A, Gorlick K, Guo J, Hillmann B, Holmes S, Holste H, Huttenhower C, Huttley GA, Janssen S, Jarmusch AK, Jiang L, Kaehler BD, Kang KB, Keefe CR, Keim P, Kelley ST, Knights D, Koester I, Kosciolek T, Kreps J, Langille MGI, Lee J, Ley R, Liu Y-X, Loftfield E, Lozupone C, Maher M, Marotz C, Martin BD, McDonald D, McIver LJ, Melnik AV, Metcalf JL, Morgan SC, Morton JT, Naimey AT, Navas-Molina JA, Nothias LF, Orchanian SB, Pearson T, Peoples SL, Petras D, Preuss ML, Pruesse E, Rasmussen LB, Rivers A, Robeson II MS, Rosenthal P, Segata N, Shaffer M, Shiffer A, Sinha R, Song SJ, Spear JR, Swafford AD, Thompson LR, Torres PJ, Trinh P, Tripathi A, Turnbaugh PJ, Ul-Hasan S, van der Hooft JJJ, Vargas F, Vázquez-Baeza Y, Vogtmann E, von Hippel M, Walters W, Wan Y, Wang M, Warren J, Weber KC, Williamson CHD, Willis AD, Zech Xu Z, Zaneveld JR, Zhang Y, Zhu Q, Knight R, Caporaso JG. 2019. Reproducible, interactive, scalable, and extensible microbiome data science using QIIME 2. Nat Biotechnol 37:852–857.

74. Callahan BJ, McMurdie PJ, Rosen MJ, Han AW, Johnson AJA, Holmes SP. 2016. DADA2: High-resolution sample inference from Illumina amplicon data. Nat Methods 13:581 –583.

75. Bokulich NA, Kaehler BD, Rideout JR, Dillon M, Bolyen E, Knight R, Huttley GA, Caporaso JG. 2018. Optimizing taxonomic classification of marker-gene amplicon sequences with QIIME 2’s q2-feature-classifier plugin. Microbiome 6:90.

76. Quast C, Pruesse E, Yilmaz P, Gerken J, Schweer T, Yarza P, Peplies J, Glöckner FO. 2013. The SILVA ribosomal RNA gene database project: Improved data processing and web-based tools. Nucleic Acids Res 41:590–596.

77. Price MN, Dehal PS, Arkin AP. 2010. FastTree 2 –Approximately Maximum-Likelihood Trees for Large Alignments. PLoS One 5:e9490.

78. R Core Team. 2008. R: A language and environment for statistical computing. R Foundation for Statistical Computing, Vienna, Austria.

79. Harrington B, Team and the D. 2005. Inkscape. Available online at: http://www.inkscape.org/.

80. QGIS Development Team. 2017. Geographic Information System. Open Source Geospatial Foundation Project. Available online at: http://qgis.osgeo.org.

81. McDougall TJ, Barker PM. 2011. Getting started with TEOS-10 and the Gibbs Seawater (GSW) Oceanographic Toolbox. SCOR/IAPSO Working Group Rep.

82. Aescht E, Büchl-Zimmermann S, Burmester A, Dänhardt-Pfeiffer S, Desel C, Hamers C, Jach G, Kässens M, Makovitzky J, Mulisch M, Nixdorf-Bergweiler B, Pütz D, Riedelsheimer B, van den Boom F, Wegerhoff R, Welsch U. 2010. Romeis Mikroskopische Technik. Springer Spektrum. Spektrum Akademischer Verlag, Heidelberg.

83. Andrews S. 2010. FastQC: a quality control tool for high throughput sequence data. Available online at: http://www.bioinformatics.babraham.ac.uk/projects/fastqc.

84. Li D, Luo R, Liu C-M, Leung C-M, Ting H-F, Sadakane K, Yamashita H, Lam T-W. 2016. MEGAHIT v1.0: A fast and scalable metagenome assembler driven by advanced methodologies and community practices. Methods 102:3–11.

85. Uritskiy GV, DiRuggiero J, Taylor J. 2018. MetaWRAP—a flexible pipeline for genome-resolved metagenomic data analysis. Microbiome 6:158.

86. Kang DD, Froula J, Egan R, Wang Z. 2015. MetaBAT, an efficient tool for accurately reconstructing single genomes from complex microbial communities. PeerJ 3:e1165.

87. Kang DD, Li F, Kirton E, Thomas A, Egan R, An H, Wang Z. 2019. MetaBAT 2: an adaptive binning algorithm for robust and efficient genome reconstruction from metagenome assemblies. PeerJ 7:e7359.

88. Wu Y-W, Simmons BA, Singer SW. 2016. MaxBin 2.0: an automated binning algorithm to recover genomes from multiple metagenomic datasets. Bioinformatics 32:605–607.

89. Bankevich A, Nurk S, Antipov D, Gurevich AA, Dvorkin M, Kulikov AS, Lesin VM, Nikolenko SI, Pham S, Prjibelski AD, Pyshkin AV, Sirotkin AV, Vyahhi N, Tesler G, Alekseyev MA, Pevzner PA. 2012. SPAdes: a new genome assembly algorithm and its applications to single-cell sequencing. J Comput Biol 19:455–477.

90. Hyatt D, Chen G-L, LoCascio PF, Land ML, Larimer FW, Hauser LJ. 2010. Prodigal: Prokaryotic gene recognition and translation initiation site identification. BMC Bioinformatics 11:119.

91. Eddy SR. 2011. Accelerated Profile HMM Searches. PLoS Comput Biol 7:e1002195.

92. Matsen FA, Kodner RB, Armbrust EV. 2010. pplacer: linear time maximum-likelihood and Bayesian phylogenetic placement of sequences onto a fixed reference tree. BMC Bioinformatics 11:538.

93. Jain C, Rodriguez-R LM, Phillippy AM, Konstantinidis KT, Aluru S. 2018. High throughput ANI analysis of 90K prokaryotic genomes reveals clear species boundaries. Nat Commun 9:5114.

94. Letunic I, Bork P. 2016. Interactive tree of life (iTOL) v3: an online tool for the display and annotation of phylogenetic and other trees. Nucleic Acids Res 44:W242–W245.

95. Segata N, Izard J, Waldron L, Gevers D, Miropolsky L, Garrett WS, Huttenhower C. 2011. Metagenomic biomarker discovery and explanation. Genome Biol 12:R60.

96. Delmont TO, Eren AM. 2018. Linking pangenomes and metagenomes: the Prochlorococcus metapangenome. PeerJ 6:e4320.

97. Eren AM, Esen ÖC, Quince C, Vineis JH, Morrison HG, Sogin ML, Delmont TO. 2015. Anvi’o: an advanced analysis and visualization platform for ‘omics data. PeerJ 3:e1319.

98. Jones P, Binns D, Chang H-Y, Fraser M, Li W, McAnulla C, McWilliam H, Maslen J, Mitchell A, Nuka G, Pesseat S, Quinn AF, Sangrador-Vegas A, Scheremetjew M, Yong S-Y, Lopez R, Hunter S. 2014. InterProScan 5: Genome-scale protein function classification. Bioinformatics 30:1236–1240.

99. Sangrador-Vegas A, Mitchell AL, Chang H-Y, Yong S-Y, Finn RD. 2016. GO annotation in InterPro: why stability does not indicate accuracy in a sea of changing annotations. Database (Oxford) 2016:baw027.

100. Elbourne LDH, Tetu SG, Hassan KA, Paulsen IT. 2017. TransportDB 2.0: a database for exploring membrane transporters in sequenced genomes from all domains of life. Nucleic Acids Res 45:D320–D324.

101. Dever M, Hebert D, Greenan BJW, Sheng J, Smith PC. 2016. Hydrography and Coastal Circulation along the Halifax Line and the Connections with the Gulf of St. Lawrence. Atmos-Ocean 54:199–217.

102. Fratantoni PS, Pickart RS. 2007. The Western North Atlantic Shelfbreak Current System in Summer. J Phys Oceanogr 37:2509–2533.

